# On the (un-)predictability of a large intragenic fitness landscape

**DOI:** 10.1101/048769

**Authors:** Claudia Bank, Sebastian Matuszewski, Ryan T. Hietpas, Jeffrey D. Jensen

**Affiliations:** Instituto Gulbenkian de Ciência, 2780-156 Oeiras, Portugal; School of Life Sciences, École Polytechnique Fédérale de Lausanne (EPFL), Lausanne, Switzerland; Swiss Institute of Bioinformatics (SIB), Lausanne, Switzerland; Eli Lilly and Company, Indianapolis, IN 46225; Department of Biochemistry & Molecular Pharmacology, University of Massachusetts Medical School, Worcester, MA 01605

**Keywords:** evolution, adaptation, epistasis, fitness landscapes

## Abstract

The study of fitness landscapes, which aims at mapping genotypes to fitness, is receiving ever-increasing attention. Novel experimental approaches combined with NGS methods enable accurate and extensive studies of the fitness effects of mutations – allowing us to test theoretical predictions and improve our understanding of the shape of the true underlying fitness landscape, and its implications for the predictability and repeatability of evolution.

Here, we present a uniquely large multi-allelic fitness landscape comprised of 640 engineered mutants that represent all possible combinations of 13 amino-acid changing mutations at six sites in the heat-shock protein Hsp90 in *Saccharomyces cerevisiae* under elevated salinity. Despite a prevalent pattern of negative epistasis in the landscape, we find that the global fitness peak is reached via four positively epistatic mutations. Combining traditional and extending recently proposed theoretical and statistical approaches, we quantify features of the global multi-allelic fitness landscape. Using subsets of the data, we demonstrate that extrapolation beyond a known part of the landscape is difficult owing to both local ruggedness and amino-acid specific epistatic hotspots, and that inference is additionally confounded by the non-random choice of mutations for experimental fitness landscapes.

**Author Summary:** The study of fitness landscapes is fundamentally concerned with understanding the relative roles of stochastic and deterministic processes in adaptive evolution. Here, the authors present a uniquely large and complete multi-allelic intragenic fitness landscape of 640 systematically engineered mutations in yeast Hsp90. Using a combination of traditional and recently proposed theoretical approaches, they study the accessibility of the global fitness peak, and the potential for predictability of the fitness landscape topography. They report local ruggedness of the landscape and the existence of epistatic hotspot mutations, which together make extrapolation and hence predictability inherently difficult, if mutation-specific information is not considered.

## Introduction

Since first proposed by Sewall Wright in 1932 [1], the idea of a fitness landscape relating genotype (or phenotype) to the reproductive success of an individual has inspired evolutionary biologists and mathematicians alike [2, 3, 4]. With the advancement of molecular and systems biology towards large and accurate data sets, it is a concept that receives increasing attention across other subfields of biology [5, 6, 7, 8, 9]. The shape of a fitness landscape carries information on the repeatability and predictability of evolution, the potential for adaptation, the importance of genetic drift, the likelihood of convergent and parallel evolution, and the degree of optimization that is (theoretically) achievable [4]. Unfortunately, the dimensionality of a complete fitness landscape of an organism – that is, a mapping of all possible combinations of mutations to their respective fitness effects – is much too high to be assessed experimentally. With the development of experimental approaches that allow for the assessment of full fitness landscapes of tens to hundreds of mutations, there is growing interest in statistics that capture the features of the landscape, and that relate an experimental landscape to theoretical landscapes of similar architecture, which have been studied extensively [10]. It is, however, unclear whether this categorization allows for an extrapolation to unknown parts of the landscape, which would be the first step towards quantifying predictability – an advance that would yield impacts far beyond the field of evolutionary biology, in particular for the clinical study of drug resistance evolution in pathogens and the development of effective vaccine and treatment strategies [8].

Existing research in this rapidly growing field comes from two sides. Firstly, different empirical landscapes have been assessed (reviewed in [4]), generally based on the combination of previously observed beneficial mutations or on the dissection of an observed adaptive walk (i.e., a combination of mutations that have been observed to be beneficial in concert). Secondly, theoretical research has proposed different landscape architectures (such as the House-of-cards, the Kauffman NK, and the Rough-Mount-Fuji model), studied their respective properties, and developed a number of statistics that characterize the landscape and quantify the expected degree of epistasis (i.e., interaction effects between mutations) [11, 12, 13, 10, 14].

The picture that emerges from these studies is mixed, reporting both smooth [15] and rugged [16, 17] landscapes with both positive epistasis (i.e., two mutations in concert are more advantageous than expected; [18]) and negative epistasis (i.e., two mutations in concert are more deleterious than expected; [19, 20]; but see [21, 22]). Current statistical approaches have been used to rank the existing landscapes by certain features [10, 14] and to assess whether they are compatible with Fisher’s Geometric Model [23]. A crucial remaining question is the extent to which the non-random choice of mutations for the experiment affects the topography of the landscape, and whether the local topography is indeed informative as to the rest of the landscape.

Here, we present an intragenic fitness landscape of 640 amino-acid changing mutations in the heat shock protein Hsp90 in *Saccharomyces cerevisiae* in a challenging environment imposed by high salinity. With all possible combinations of 13 mutations of various fitness effects at 6 positions, the presented landscape is not only uniquely large but also distinguishes itself from previously published work regarding several other experimental features – namely, by its systematic and controlled experimental setup using engineered mutations of various selective effects, and by considering multiple alleles simultaneously. We begin by describing the landscape and identifying the global peak, which is reached through a highly positively epistatic combination of four mutations. Based on a variety of implemented statistical measures and models, we describe the accessibility of the peak, the pattern of epistasis, and the topography of the landscape. In order to accommodate our data, we extend several previously used models and statistics to the multi-allelic case. Using subsets of the landscape, we discuss the predictive potential of such modeling and the problem of selecting non-random mutations when attempting to quantify local landscapes in order to extrapolate global features.

## Material and Methods

Here, we briefly outline the materials and methods used. A more detailed treatment of the theoretical work is presented in the Supporting Information.

### Data generation

Codon substitution libraries consisting of 640 combinations (single, double, triple and quadruple mutants) of 13 previously isolated individual mutants within the 582-590 region of yeast Hsp90 were generated from optimized cassette ligation strategies as previously described in [24] and cloned into the p417GPD plasmid that constitutively expresses Hsp90.

Constitutively expressed libraries of Hsp90 mutation combinations were introduced into the *S. cerevisiae* shutoff strain DBY288 (can1-100 ade2-1 his3-11,15 leu2-3,12 trp1-1 ura3-1 hsp82: :leu2 hsc82: :leu2 ho: : :pgals-hsp82-his3) using the lithium acetate method [25]. Following transformation the library was amplified for 12 hours at 30deg C under nonselective conditions using galactose (Gal) medium with 100 *μ*g/mL ampicillin (1.7 g yeast nitrogen base without amino acids, 5 g ammonium sulfate, 0.1 g aspartic acid, 0.02 g arginine, 0.03 g valine, 0.1 g glutamic acid, 0.4 g serine, 0.2 g threonine, 0.03 g isoleucine, 0.05 g phenylalanine, 0.03 g tyrosine, 0.04 g adenine hemisulfate, 0.02 g methionine, 0.1 g leucine, 0.03 g lysine, 0.01 g uracil per liter with 1% raffinose and 1% galactose). After amplification the library culture was transferred to selective medium similar to Gal medium but raffinose and galactose are replaced with 2% dextrose. The culture was grown for 8 hours at 30deg C to allow shutoff of the wild-type copy of Hsp90 and then shifted to selective medium containing 0.5M NaCl for 12 generations. Samples were taken at specific time points and stored at −80degC.

Yeast lysis, DNA isolation and preparation for Illumina sequencing were performed as previously described [26]. Sequencing was performed by Elim Biopharmaceuticals, Inc and produced ≈30 million reads of 99% confidence at each read position based on PHRED scoring [27, 28]. Analysis of sequencing data was performed as previously described [29].

### Estimation of growth rates

Individual growth rates were estimated according to the approach described by [20] using a Bayesian Monte Carlo Markov Chain (MCMC) approach proposed in [30]. Nucleotide sequences coding for the same amino acid sequence were interpreted as replicates with equal growth rates. The resulting MCMC output consisted of 10,000 posterior estimates for each amino acid mutation corresponding to an average effective samples size of 7,419 (minimum 725). Convergence was assessed using the Hellinger distance approach [31] combined with visual inspection of the resulting trace files.

### Adaptive walks

In the strong selection weak mutation (SSWM) limit [32], adaptation can be modeled as a Markov process only consisting of subsequent fitness-increasing one-step substitutions that continue until an optimum is reached (so-called adaptive walks). This process is characterized by an absorbing Markov chain with a total of *n* different states (i.e., mutants), consisting of *k* absorbing (i.e., optima) and *n* – *k* transient states (i.e., non-optima). Defining *w*(*g*) as the fitness of genotype *g*, and *g*_[*i*]_ as the genotype *g* carrying a mutant allele at locus i, the selection coefficient is denoted by *S*_*j*_(*g*) = *w*(*g*_[*j*]_) – *w*(*g*), such that the transition probabilities *p*_*g,g*[*i*]_ for going from any mutant *g* to any mutant *g*_[*i*]_ are given by the selection coefficient normalized by the sum over all adaptive, one-mutant neighbors of the current genotype *g*. If *g* is a (local) optimum, *p*_*g*,*g*_ = 1. Putting the transition matrix **P** in its canonical form and computing the fundamental matrix, then allows to determine the expectation and the variance in the number of steps before reaching *any* optimum, and to calculate the probability to reach optimum *g* when starting from genotype *g*′ [33]. Robustness of the results and the influence of specific mutations were assessed by deleting the corresponding columns and rows in P (i.e., by essentially treating the corresponding mutation as unobserved), and re-calculating and comparing all statistics to those obtained from the full data set.

### Correlation of fitness effects of mutations

Strength and type of epistasis was assessed by calculating the correlation of fitness effects of mutations *γ* [14], which quantifies how the selective effect of a focal mutation is altered when put onto a different genetic background, averaged over all genotypes of the fitness landscape. Extending recent theory [14], we calculated the matrix of epistatic effects between different pairs of alleles (*A*_*i*_,*B*_*i*_) and (*A*_*j*_,*B*_*j*_) termed 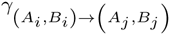 (eq. S1_8), the vector of epistatic effects between a specific pair of alleles (*A*_*i*_,*B*_*i*_) on all other pairs of alleles 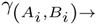 (eq. S1_9), the vector of epistatic effects between all pairs of alleles on a specific allele pair (*A*_*j*_,*B*_*j*_) termed 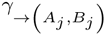 (eq. S1_12), and the decay of correlation of fitness effects *γ*_*d*_ (eq. S1_15) with Hamming distance *d* averaged over all genotypes *g* of the fitness landscape.

### Fraction of epistasis

Following [34] and [35], we quantified whether specific pairs of alleles between two loci interact epistatically, and if so whether these display magnitude epistasis (i.e., fitness effects are nonadditive, but fitness increases with the number of mutations), sign epistasis (i.e., one of the two mutations considered has an opposite effect in both backgrounds) or reciprocal sign epistasis (i.e., if both mutations show sign epistasis). In particular, we calculated the type of epistatic interaction between mutations *g*_[*i*]_ and *g*_[*j*]_ (with *i* ≠ *j*) with respect to a given reference genotype *g* over the entire fitness landscape. There was no epistatic interaction if 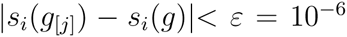, magnitude epistasis if 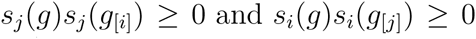, reciprocal sign epistasis if *s*_*j*_ (*g*)*s_j_* (*g*_[*i*]_) < 0 and *s*_*i*_(*g*)*s*_*i*_(*g*_[*j*]_) < 0, and sign epistasis in all other cases [36].

### Roughness-to-slope ratio

Following [11], we calculated the roughness-to-slope ratio *ρ* by fitting the fitness landscape to a multidimensional linear model using the least-squares method. The slope of the linear model corresponds to the average additive fitness effect [10, 23], whereas the roughness is given by the variance of the residuals. Generally, the better the linear model fit, the smaller the variance in residuals such that the roughness-to-slope ratio approaches 0 in a perfectly additive model. Conversely, a very rugged fitness landscape would have a large residual variance and, thus, a very large roughness-to-slope ratio (as in the House-of-Cards model). In addition, we calculated a test statistic *Z*_*ρ*_ by randomly shuffling fitness values in the sequence space to evaluate the statistical significance of the obtained roughness-to-slope ratio from the data set *ρ*_*data*_ given by 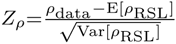.

## Results and Discussion

We used the EMPIRIC approach [24, 26] to assess the growth rate of 640 mutants in yeast Hsp90 (see Materials and Methods). Based on previous screenings of fitness effects in different environments [29] and on different genetic backgrounds [20], and on expectations of their biophysical role, 13 amino-acid changing point mutations at 6 sites were chosen for the fitness landscape presented here (Fig. 1). The fitness landscape was created by assessing the growth rate associated with each individual mutation on the parental background, and all possible mutational combinations. A previously described MCMC approach was used to assess fitness and credibility intervals ([30]; see Materials and Methods).

**Figure 1.**
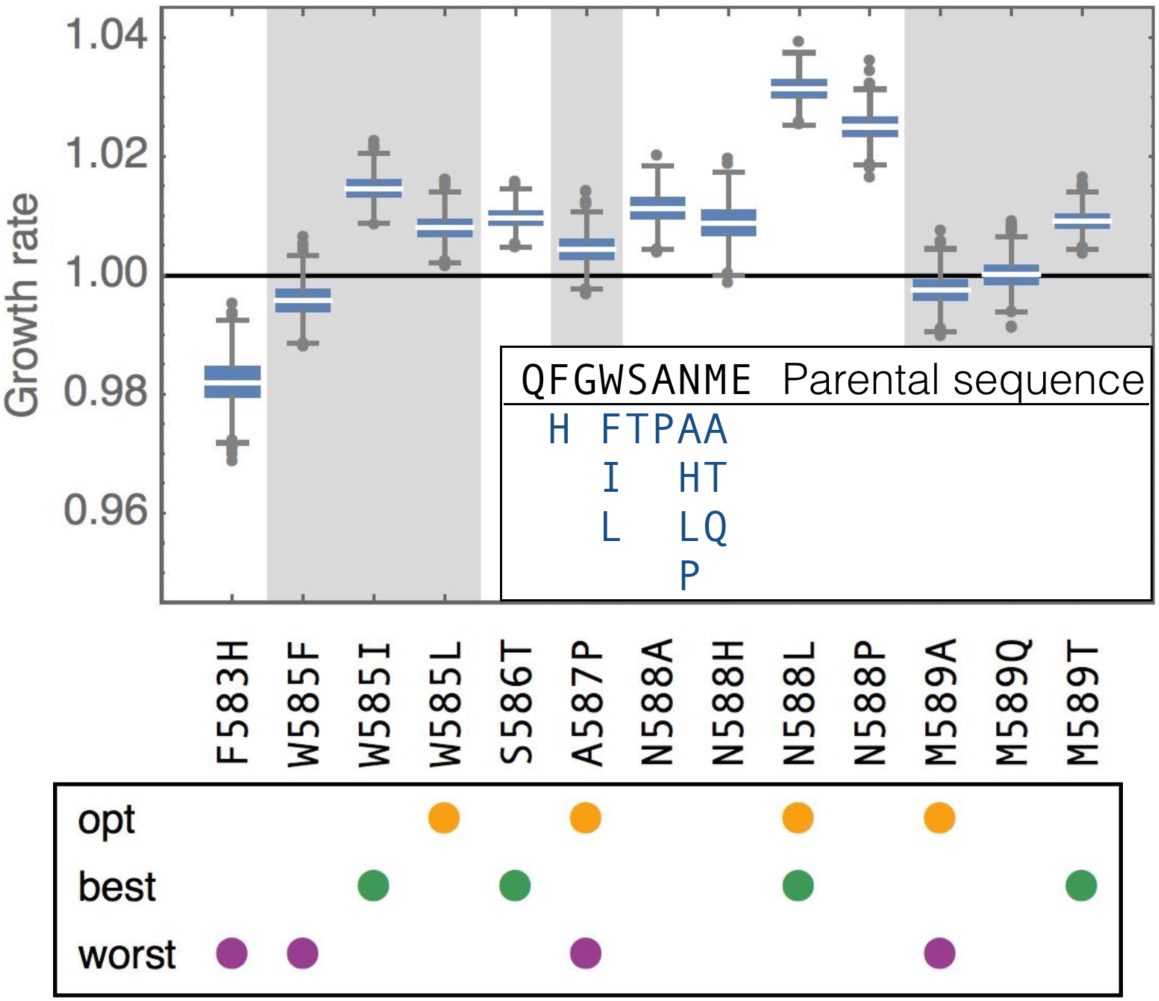
Individual amino-acid substitutions and their effect on the parental background in elevated salinity, obtained from 1000 samples from the posterior distribution of growth rates. Boxes represent the interquartile range (i.e., the 50% C.I.), whiskers extend to the highest/lowest data point within the box ± 1.5 times the interquartile range, and circles represent outliers; gray and white background shading alternates by amino acid position. The box below indicates with colored dots which mutations are involved in the focal landscapes discussed throughout the main text: the four mutations leading to the global optimum (“opt”), the four individually most beneficial mutations (“best”) and the four mutations with the individually lowest growth rates (“worst”). Inset: parental sequence at positions 582-590 and assessed amino acids by position.

### The fitness landscape and its global peak

Figure 2 presents the resulting fitness landscape, with each mutant represented based on its Hamming distance from the parental genotype and its median estimated growth rate. Lines connect single-step substitutions, with vertical lines occurring when there are multiple mutations at the same position (Fig. 1). With increasing Hamming distance from the parental type, many mutational combinations become strongly deleterious. This indicates strong negative epistasis between the substitutions that, as single steps on the background of the parental type, have small effects. This pattern is consistent with Fisher’s Geometric Model [37] when combinations of individually beneficial or small-effect mutations overshoot the optimum, and with classical arguments predicting negative epistasis based on mutational load [38]. It is also intuitively comprehensible on the protein level, where the accumulation of too many mutations is likely to destabilize the protein and render it dysfunctional [39].

**Figure 2.**
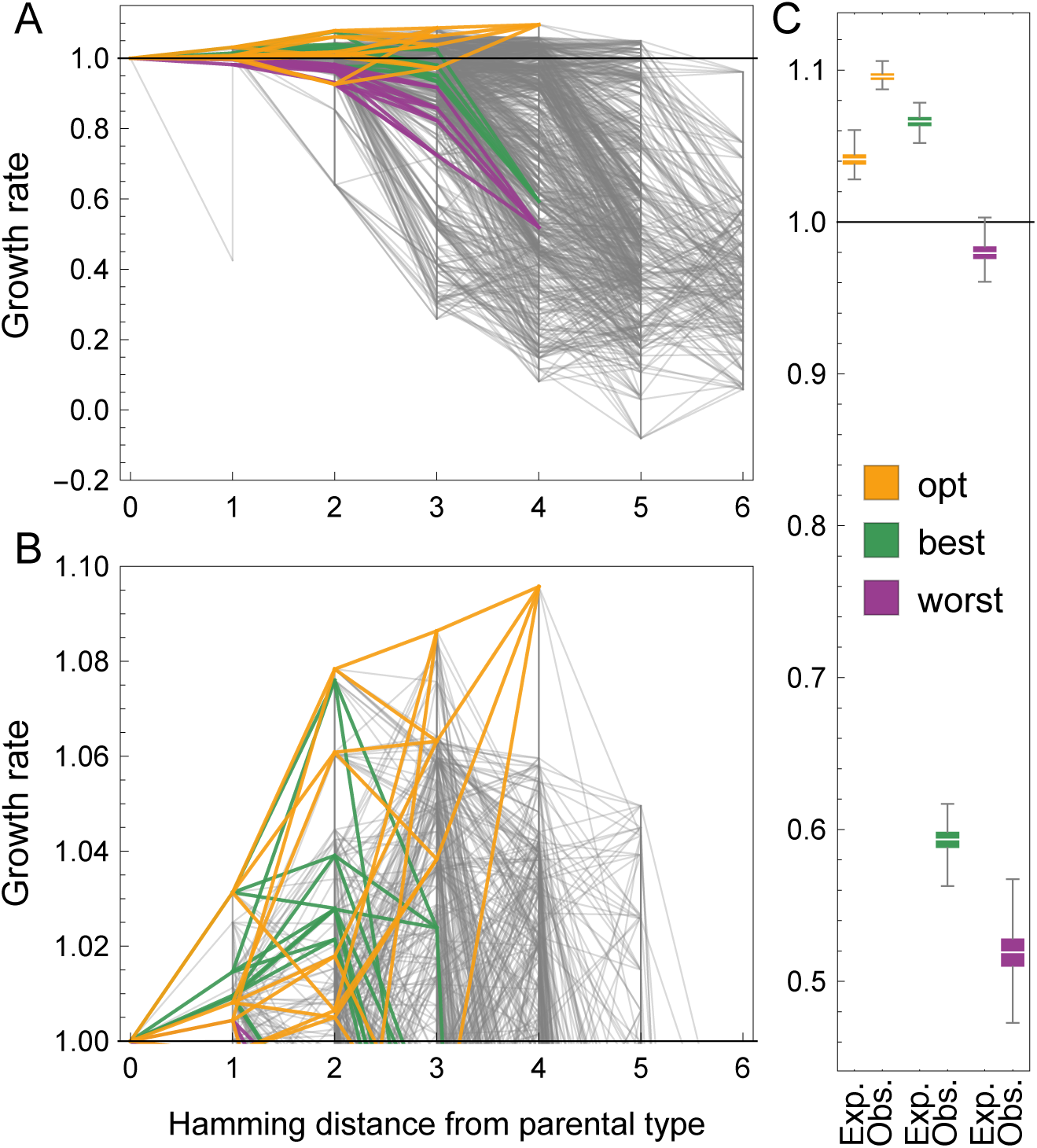
(A) Empirical fitness landscape by mutational distance to parental type. Each line represents a single substitution. Vertical lines appear when multiple alleles have been screened at the same position. There is a global pattern of negative epistasis. The three focal landscapes are highlighted. (B) Close-up on the beneficial portion of the landscape. (C) Expected (“Exp.”) versus observed (“Obs.”) fitness for the focal landscapes, obtained from 1000 posterior samples. We observe strong positive epistasis in the landscape that contains the global optimum, whereas the other two are dominated by negative epistasis. In all panels, the y axis depicts growth rate as a proxy for fitness.

The global peak of the fitness landscape is located 4 mutational steps away from the parental type (Fig. 2B), with 98% of posterior samples identifying the peak. The fitness advantage of the global peak reaches nearly 10% over the parental type, and is consistent between replicates (see Materials and Methods; Fig. S3_1). Though perhaps surprising given the degree of conservation of the studied genomic region ([24], their Fig. S5), it is important to note that these fitness effects are measured under highly artificial experimental conditions including high salinity, which are unlikely to represent a natural environment of yeast. The effects of the individual mutations comprising the peak in a previous experiment without added NaCl were −0.04135, −0.01876, −0.03816, −0.02115 for mutations W585L, A587P, N588L and M589A, respectively, emphasizing the potential cost of adaptation associated with the increased salinity environment (data from [20]; see also [29]).

Curiously, the global peak is not reached by combining the most beneficial single-step mutations, but via a highly synergistic combination of one beneficial and three ‘neutral’ mutations (i.e., mutations that are individually indistinguishable from the parental type in terms of growth rate). In fact, each one of the five beneficial local optima shows a similar signature of positive epistasis (Fig. S3_2). Figure 2C demonstrates that a combination of the four mutations involved in the global peak (termed “opt”) predicts only a 4% fitness advantage. Furthermore, even a combination of the four individually most beneficial single-step mutations in the data set (“best”; considering at most one mutation per position) only predicts a benefit of 6%. Notably, the actual combination of these four mutations is highly deleterious and thus exhibits strong negative epistasis. Although negative epistasis between beneficial mutations during adaptation has been reported more frequently, positive epistasis has also been observed occasionally [18, 40], particularly in the context of compensatory evolution. In fact, negative epistasis between beneficial mutations and positive epistasis between neutral mutations has been predicted by de Visser *et al.* [41]. Furthermore, our results support the pattern recently found in the gene underlying the antibiotic resistance enzyme TEM-1 beta-lactamase in *E. coli*, showing that large-effect mutations interact more strongly than small-effect mutations such that the fitness landscape of large-effect mutations tends to be more rugged than the landscape of small-effect mutations [13], and that mutations that were selected for their combined beneficial effect tend to interact synergistically, whereas mutations selected for their individual effects interact antagonistically [13, 4].

### Adaptive walks on the fitness landscape

Next, we studied the empirical fitness landscape within a framework recently proposed by Draghi and Plotkin [42]. Given the empirical landscape, we simulated adaptive walks and studied the accessibility of the six observed local optima. In addition, we evaluated the length of adaptive walks starting from any mutant in the landscape, until an optimum is reached. In the strong selection weak mutation limit [43], we can express the resulting dynamics as an absorbing Markov chain, where local optima correspond to the absorbing states, and in which the transition probabilities correspond to the relative fitness increases attainable by the neighboring mutations (see Materials and Methods). This allowed us to derive analytical solutions for the mean and variance of the number of steps to reach a fitness optimum (see extended Materials and Methods), and the probability to reach a particular optimum starting from any given mutant in the landscape (Figs. 3, S3_3, S3_4).

Using this framework, we find that the global optimum can be reached with non-zero probability from almost 95% of starting points in the landscape, and is reached with high probability from a majority of starting points - indicating high accessibility of the global optimum (Fig. 3). The picture changes when restricting the analysis to adaptive walks initiating from the parental type (Figs. 3, S3_3, S3_4). Here, although 73% of all edges and 78% of all vertices are included in an adaptive walk to the global optimum, it is reached with only 26% probability. A local optimum two substitutions away from the parental type (Fig. 3C) is reached with a much higher probability of 47%. Hence, adaptation on the studied landscape is likely to stall at a sub-optimal fitness peak. This indicates that the local and global landscape pattern may be quite different, an observation that is confirmed and discussed in more detail below. In line with the existence of multiple local fitness peaks, we find that pairs of alleles at different loci show pervasive sign (30%) and reciprocal sign (8%) epistasis [34], whereas the remaining 62% are attributed to magnitude epistasis (i.e., there is no purely additive interaction between alleles; for a discussion of the contribution of experimental error see Fig. S3_5).

**Figure 3.**
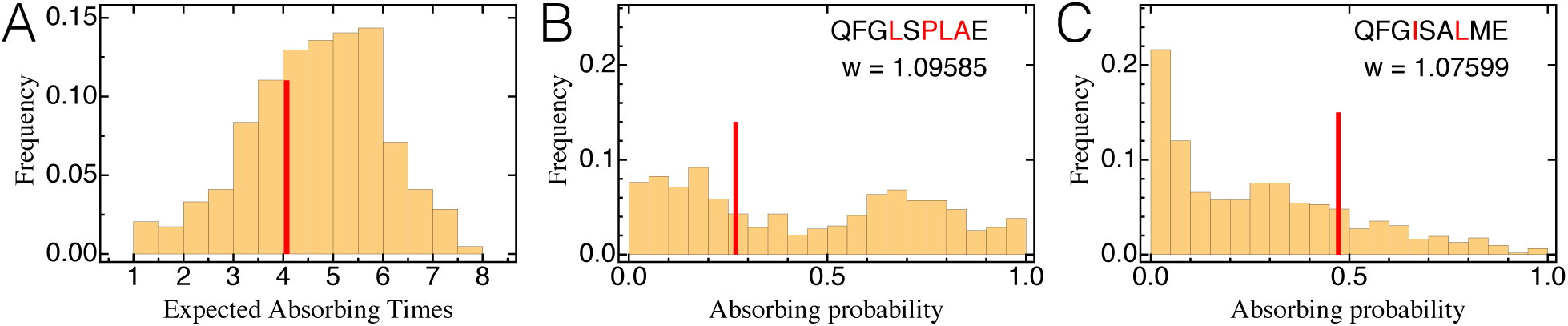
(A) Distribution of average lengths of adaptive walks starting from any type in the full landscape (i.e., absorbing times of the Markov chain). The red line indicates mean path length for adaptive walks from the parental genotype. (B) & (C): Distribution of absorbing probabilities, that is, the probability to reach a specific optimum starting from a given genotype computed for all genotypes in the data set. The red line corresponds to the respective probability when starting from the parental sequence. The global optimum (panel B) is in general reached with a very high probability, but there are starting points from which it is poorly accessible.

### Epistasis measures and the topography of the fitness landscape

Next, we considered the global topography of the fitness landscape. Various measures of epistasis and ruggedness have been proposed, most of them correlated and hence capturing similar features of the landscape [10]. However, drawing conclusions has proven difficult because the studied landscapes were created according to different criteria. Furthermore, the majority of published complete landscapes are too small to be divided into subsets [but see 10, 12], preventing tests for the consistency and hence the predictive potential of landscape statistics. The landscape studied here provides us with this opportunity. Moreover, because multiple alleles at the same site are contained within the landscape, we may study whether changes in the shape of the landscape are site- or amino-acid specific.

We computed various landscape statistics (roughness-to-slope ratio, fraction of epistasis, and the recently proposed gamma statistics; see SI 1) [11, 10, 14], and compared them to expectations from theoretical landscape models (NK, Rough-Mount-Fuji (RMF), House-of-Cards (HoC), Egg-Box landscapes; for brief definitions of these see SI 2). Whenever necessary, we provide an analytical extension of the used statistic to the case of multi-allelic landscapes (see Materials and Methods). To assess consistency and predictive potential, we computed the whole set of statistics for (1) all landscapes in which one amino acid was completely removed from the landscape (a cross-validation approach [44], subsequently referred to as the ‘drop-one’ approach), (2) all possible 360 di-allelic sub-landscapes, and (3) for all 1,570 di-allelic 4-step landscapes containing the parental genotype, highlighting as special examples the three focal landscapes discussed.

We find that the general topography of the fitness landscape resembles that of a RMF landscape with intermediate ruggedness, which is characterized by a mixture of a random HoC component and an additive component (Fig. 4A,B). Whereas the whole set of landscape statistics supports this topography and our conclusions, the gamma statistics measuring landscape-averaged correlations in fitness effects, recently proposed by [14], proved to be particularly illustrative. We will therefore focus on these in the main text; we refer to the Supplementary Material for additional results (e.g., measurement uncertainty and adaptive walks).

**Figure 4.**
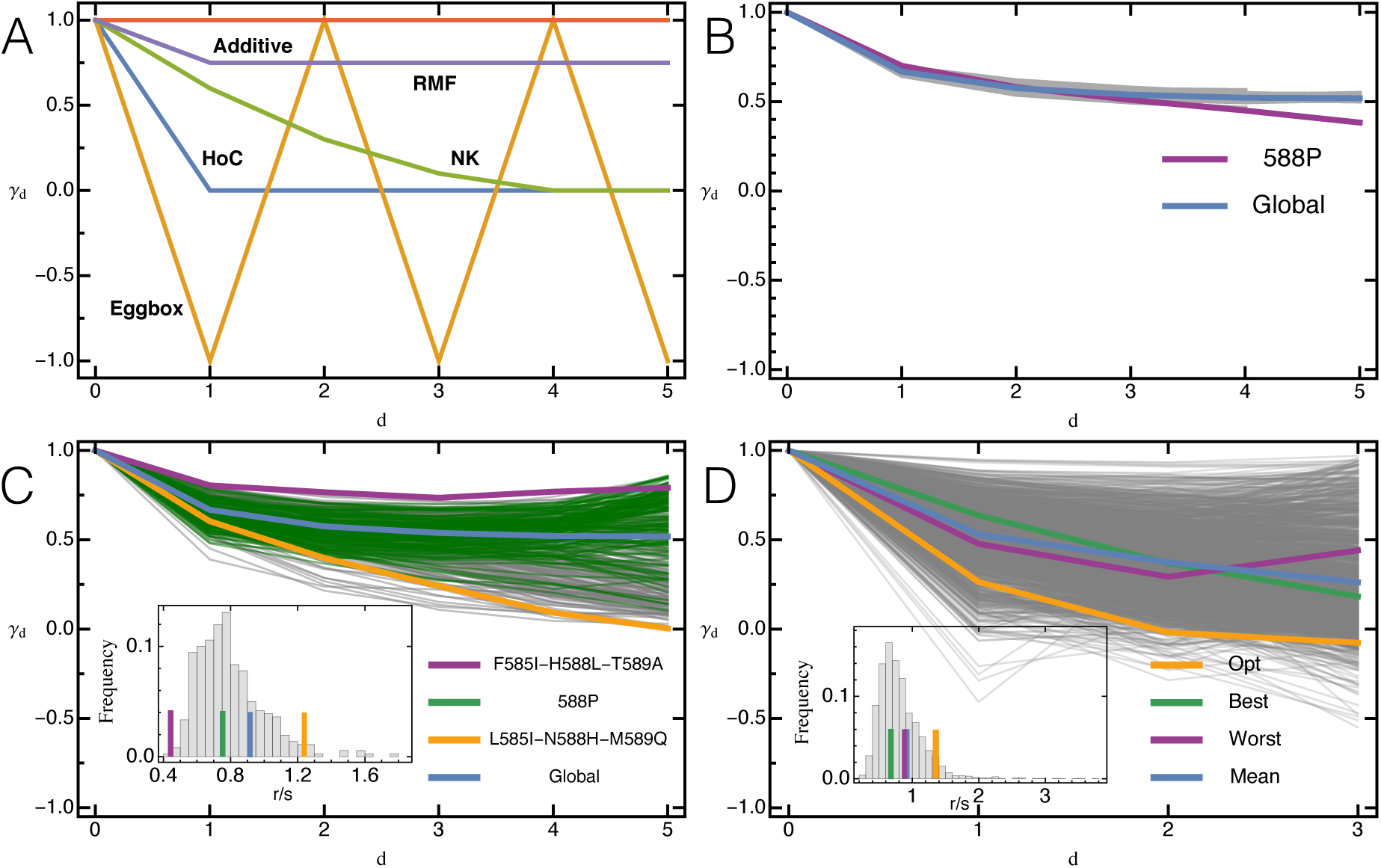
(A) Expected pattern of landscape-wide epistasis measure *γ*_*d*_ (eq. S1_15) with mutational distance for theoretical fitness landscapes with 6 loci from [14]. (B) Observed decay of *γ*_*d*_ with mutational distance under the drop-one approach is quite homogenous, except when hotspot mutation 588P is removed. 95% C.I. are contained within the lines for the global landscape. (C) Observed decay of *γ*_*d*_ for all di-allelic 6-locus sub-landscapes. Depending on the underlying mutations, *γ*_*d*_ is vastly different suggesting qualitative differences in the topography of the underlying fitness landscape and in the extent of additivity in the landscape. Three focal landscapes representative of different types, and *γ*_*d*_ for the full landscape have been highlighted. (D) Observed decay of *γ*_*d*_ for all di-allelic 4-locus sub-landscapes containing the parental type, indicative of locally different landscape topographies. Highlighted are the three focal landscapes. Insets: Histograms of the roughness-to-slope ratio *r*/*s* for the respective subset, with horizontal lines indicating values for highlighted landscapes (*r*/*s* for global landscape (blue) is significantly different from HoC expectation, *p =* 0.01). Similar to *γ*_*d*_, there is a huge variation in *r*/*s* for subparts of the fitness landscape, especially when considering the 4-locus subsets.

### Predictive potential of landscape statistics

When computed based on the whole landscape and on a drop-one approach, the landscape appears quite homogeneous, and the gamma statistics show relatively little epistasis (Fig. 4B, 5B). On first sight, this contradicts our earlier statement of strong negative and positive epistasis but can be understood given the different definitions of the epistasis measures used: Above, we have measured epistasis based on the deviation from the multiplicative combination of the single-step fitness effects of mutations on the parental background. As these effects were small, epistasis was strong in comparison. Conversely, the gamma measure is independent of a reference genotype and captures the fitness decay with a growing number of substitutions as a dominant and quite additive component of the landscape.

Only mutation 588P has a pronounced effect on the global landscape statistics, and seems to act as an epistatic hotspot by making a majority of subsequent mutations (of individually small effect) on its background strongly deleterious (clearly visible in Fig. 4B, 5C). This can be explained by looking at the biophysical properties of this mutation. In wild-type Hsp90, amino acid 588N is oriented away from solvent and forms hydrogen bond interactions with neighboring amino acids [24]. Proline lacks an amide proton, which inhibits hydrogen bond interactions. As a result, substituting 588N with a proline could disrupt hydrogen bond interactions with residues that may be involved in main chain hydrogen bonding and destabilize the protein. In addition, the pyrrolidine ring of proline is extremely rigid and can constrain the main chain, which may restrict the conformation of the residue preceding it in the protein sequence [45].

The variation between inferred landscape topographies increases dramatically for the 360 di-allelic 6-locus sub-landscapes (Fig.4C). Whereas they are still largely compatible with an RMF landscape, the decay of landscape-wide epistasis with mutational distance (as measured by *γ*_*d*_) shows a large variance, suggesting large differences in the degree of additivity. Interestingly, various sub-landscapes, typically carrying mutation *588P, show a relaxation of epistatic constraint with increasing mutational distance (i.e., increasing *γ*_*d*_) that is not captured by any of the proposed theoretical fitness landscape models, suggestive of systematic compensatory interactions (but see the eggbox model for an explicit example featuring non-monotonicity in *γ*_*d*_). The variation in the shape of the fitness (sub-)landscapes is also reflected in the corresponding roughness-to-slope ratio (inset of Fig. 4C-D), further emphasizing inhomogeneity of the fitness landscape with local epistatic hotspots.

Finally, the 1,570 di-allelic 4-locus landscapes containing the parental genotype, though highly correlated genetically, reflect a variety of possible landscape topographies (Fig.4D), ranging from almost additive to egg-box shapes, accompanied by an extensive range of roughness-to-slope ratios. The three focal landscapes discussed above are not strongly different compared with the overall variation; yet show diverse patterns of epistasis between substitutions (Fig. 5).

**Figure 5.**
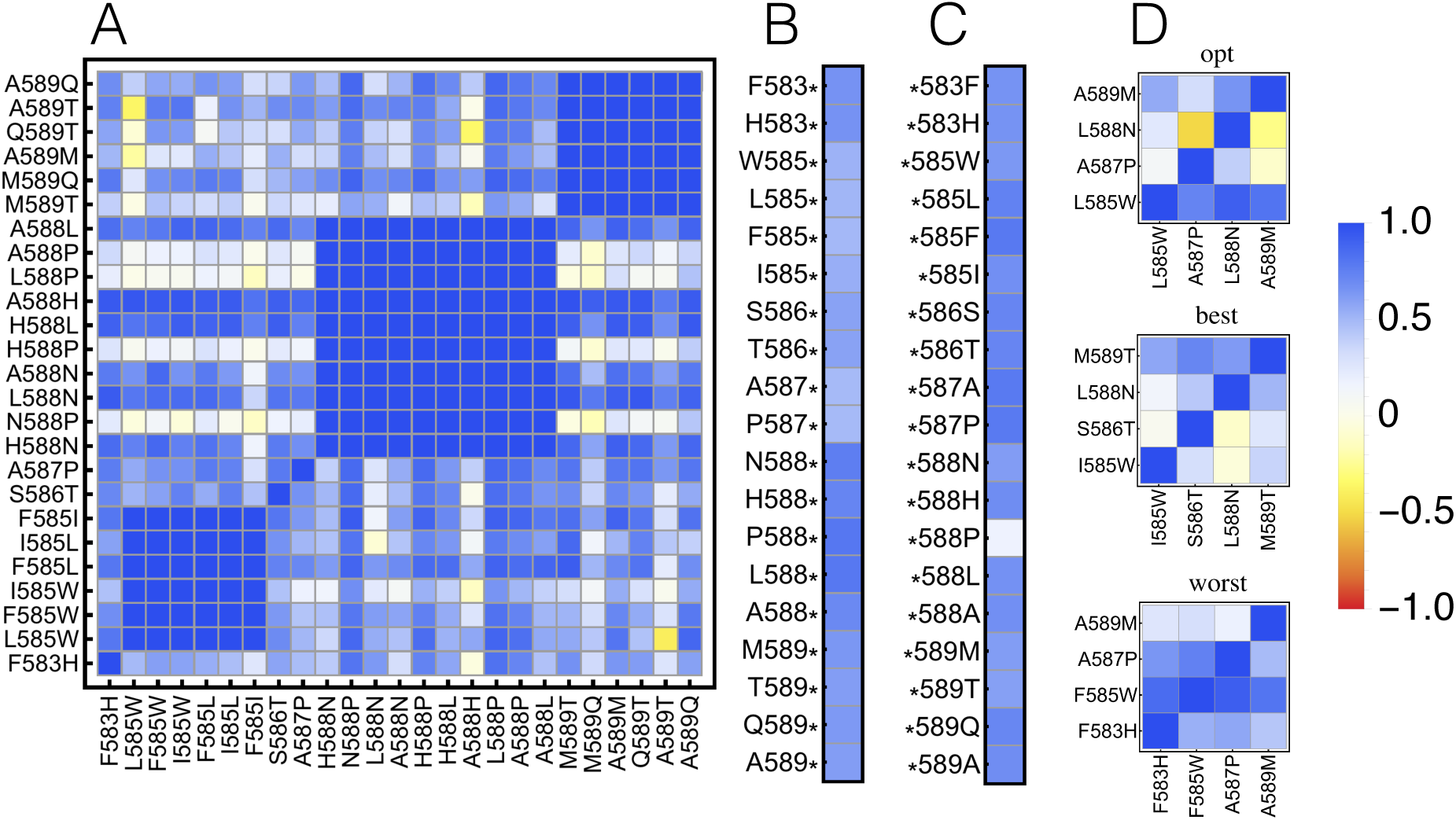
(A) Epistasis measure 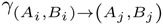 (eq. S1_8) between any two substitutions, averaged across the entire landscape. The majority of interactions are small to moderate (blue). Parts of the fitness landscape show highly localized and mutation-specific epistasis, ranging from strong magnitude epistasis (white) to sign and reciprocal sign epistasis (yellow). (B) The average epistatic effect *γA*_*i*_→(eq. S1_10) of a mutation occurring on any background is always small. (C) The average epistatic effect *γ*→*A*_*j*_ (eq. S1_13) of a background on any new mutation is usually small, except for background *588P, which shows a strong magnitude effect. (D) Locus-specific gamma for the 4 mutations leading to the global optimum (top), the 4 largest single effect mutations (middle), and the single effect mutations with the lowest fitness (bottom). The “opt” landscape exhibits strong sign epistasis between locus 587 and 588, and 588 and 589. Also the “best” landscape exhibits pervasive epistasis with sign epistasis between locus 588 and locus 585 and 586, respectively. We observe almost no epistasis in the “worst” landscape.

Thus, predicting fitness landscapes is difficult indeed. Extrapolation of the landscape, even across only a single mutation, may fail due to the existence of local epistatic hotspot mutations. While the integration of biophysical properties into landscape models is an important step forward [e.g. 46], we demonstrate that such models need to be mutation-specific. Considering a site-specific model (e.g., BLOSUM matrix; [47]) is not sufficient. Newer models such as DeepAlign may provide the opportunity to allow integration of mutation-specific effects via aligning two protein structures based on spatial proximity of equivalent residues, evolutionary relationship and hydrogen bonding similarity [48].

## Conclusion

Originally introduced as a metaphor to describe adaptive evolution, fitness landscapes promise to become a powerful tool in biology to address complex questions regarding the predictability of evolution and the prevalence of epistasis within and between genomic regions. Due to their high-dimensional nature, however, the ability to extrapolate will be paramount to progress in this area, and the optimal quantitative and qualitative approaches to achieve this goal are yet to be determined.

Here, we have taken an important step towards addressing this question via the creation and analysis of a landscape comprising 640 engineered mutants of the Hsp90 protein in yeast. The unprecedented size of the fitness landscape along with the multi-allelic nature allows us to test whether global features could be extrapolated from subsets of the data. Although the global pattern indicates a rather homogeneous landscape, smaller sub-landscapes are a poor predictor of the overall global pattern because of ‘epistatic hotspots’.

In combination, our results highlight the inherent difficulty imposed by the duality of epistasis for predicting evolution. In the absence of epistasis (i.e., in a purely additive landscape) evolution is globally highly predictable as the population will eventually reach the single fitness optimum, but the path taken is locally entirely unpredictable. Conversely, in the presence of (sign and reciprocal sign) epistasis evolution is globally unpredictable, as there are multiple optima and the probability to reach any one of them depends strongly on the starting genotype. At the same time, evolution may become locally predictable with the population following obligatory adaptive paths that are a direct result of the creation of fitness valleys owing to epistatic interactions.

The empirical fitness landscape studied here appears to be intermediate between these extremes. Although the global peak is within reach from almost any starting point, there is a local optimum that will be reached with appreciable probability, particular when starting from the parental genotype. From a practical standpoint, these results thus highlight the danger inherent to the common practice of constructing fitness landscapes from ascertained mutational combinations. However, this work also suggests that one promising way forward for increasing predictive power will be the utilization of multiple small landscapes used to gather information about the properties of individual mutations, combined with the integration of site-specific biophysical properties.

## Acknowledgements

This project was funded by grants from the Swiss National Science Foundation (FNS) and a European Research Council (ERC) Starting Grant to JDJ. Computations were performed at the Vital-IT (http://www.vital-it.ch) Center for high-performance computing of the Swiss Institute of Bioinformatics (SIB). We thank Dan Bolon, Brian Charlesworth, Pamela Cote, Inês Fragata, and Hermina Ghenu for helpful comments and discussion.

## Supporting Information 1: Extended Materials and Methods

### Adaptive walks

Under the strong selection weak mutation (SSWM) limit [43], adaptation follows an absorbing Markovian process characterized by a series of fitness-increasing substitutions along one-mutant neighbours until reaching a fitness optimum (forming a so-called adaptive walk), with a total of *n* different states (i.e., mutants), consisting of *k* absorbing (i.e., optima) and *n* – *k* transient states (i.e., non-optima).

Defining *w*(*g*) as the fitness of genotype *g*, and *g*_[*j*]_ as the genotype *g* carrying a mutant allele at locus *j*, the selection coefficient is denoted by

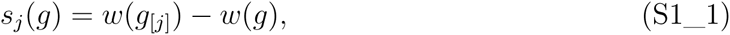

such that the transition probabilities for going from any given genotype *g* to any mutant genotype 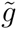 are given by the transition matrix **P**

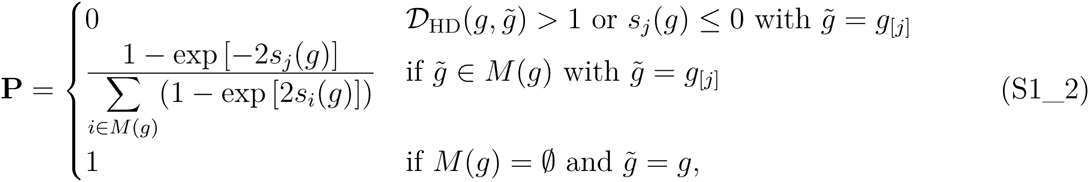

where *D*_HD_ 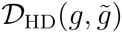 marks the Hamming distance between two genotypes *g* and 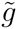 (i.e., the sum of pairwise allele differences across all loci), and 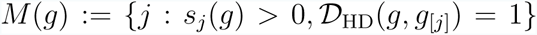 denotes the set of all adaptive, one-mutant neighbours of the current genotype *g*. Note that when considering more than two alleles per locus, *g*_[*j*]_ (and accordingly *S*_*j*_(*g*)) are not uniquely determined and – apart from the focal genotype *g* – depend on the (mutant) allele found at locus *j*. However, since both entities become explicit once 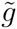 is specified (and for keeping notation simple), we prefer to define a single selection coefficient (i.e., eq. S1_1) only, that is used consistently in all our approaches. Furthermore note, that transition probabilities are given by the classical fixation probability of a given genotype [49, 50] renormalized over the fixation probability of all beneficial genotypes in a one-mutant neighbourhood of the current genotype *g*. [51] recently used an even more general form of **P** which also took into account differences in mutation rates and also allowed for the fixation of slightly deleterious and neutral mutations. However, under the assumption that there is no mutational bias (i.e., all single-step mutants are equally likely to occur) and that selection is strong as compared to mutation (i.e., the SSWM regime), both definitions are identical.

The canonical form of **P** can then be obtained by permutation, such that

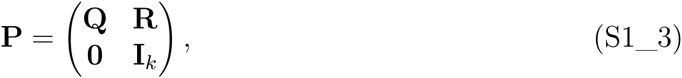

where **Q** is a (*n* – *k*) × (*n* – *k*) matrix which contains the transition probabilities between transient states; **R** is a (*n* – *k*) × *k* matrix which gives the transition probabilities from any transient to any absorbing state; **0** is the *k* × (*n* – *k*) zero matrix; and **I**_*k*_ is the *k* × *k* identity matrix [33].

Using the above representation, all basic properties of the absorbing Markov chain can be calculated from the fundamental matrix

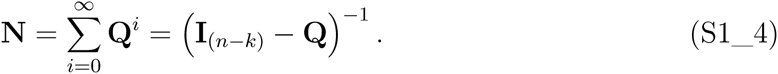

In particular, the expected number of steps before absorption (i.e., the expected number of steps on the fitness landscape before reaching any optimum) is given by

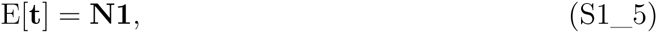

where **1** is a column vector of length (*n* – *k*) with all entries being 1, and the *i*^*th*^ entry of E[t] gives the expected number of steps when starting from state (mutant) *i*.

Similarly, the variance in the number of steps before being absorbed can be computed as

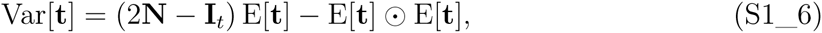

where ⊙ denotes the Hadamard product.

Finally, the probability of being absorbed in state *j* when starting from transient state *i* (i.e., reaching optimum *j* when the initial genotype is *i*), is given by the (*n* – *k*) × *k*-matrix

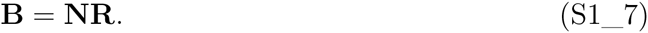

Thus, these methods give an easy and computationally fast way of quantifying and predicting adaptive walks on fitness landscapes. Furthermore, the robustness of these results and the influence of particular mutants can be assessed by deleting the *i*^*th*^ column and row of **P** – i.e., by essentially treating mutant *i* as unobserved –, recalculating the above statistics and comparing these to the statistics obtained from the full data set. Similarly, entire mutations (i.e., amino acids) can be left out to assess their relative effect on the fitness landscape and the generality of our analysis (and the statistics used).

### Measuring epistasis

We applied different metrics for quantifying epistasis over the entire fitness landscape as well as for particular mutations and assessed their consistency in capturing the strength of gene×gene interactions. In particular, we follow the definition of epistasis by [52] (originally termed epistacy), as the deviation from additivity when combining two genetic effects which is measured by the difference in log-fitness between the effects of the double mutant and the single mutant relative to the wild-type fitness.

### Correlation of fitness effects of mutations: *γ*

The first measure has recently been introduced by [14] and is defined as the single-step correlation of fitness effects for mutations between neighbouring genotypes. It quantifies how the selective effect of a focal mutation is altered when it occurs in a different genetic background averaged over all genotypes of the fitness landscape. Geometrically, *γ* measures the correlation between slopes (with respect to genotype-fitness hypercube) of the same mutation put into different genetic backgrounds. Thus, if the fitness effect of a mutation is independent of its genetic background (i.e., if there is no epistasis), the correlation in slopes will be perfect (*γ* = 1), whereas it will be zero if the fitness slopes of each genotype are independent of the fitnesses of other genotypes (as in the House-of-Cards model; 53). Depending on the scale *γ* can either be used to quantify the strength of gene×gene interactions between specific mutations or as an overall measure for the entire fitness landscape. However, in its original form *γ* was defined for di-allelic data only and thus needs to be extended by considering pairs of specific alleles at different loci [14].

Let *A_i_* = {1_*i*_, 2_*i*_,…, *m_i_*} denote the set of different alleles present at locus *i* for all polymorphic loci *i* ∈ {1, 2,…, *n*} such that 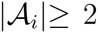 for all loci *i*. Further, let 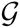 be the set of all genotypes that can be formed by combining all alleles such that the total number of genotype is 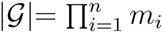.

Then the matrix of epistatic effects between loci *i* and *j* carrying alleles *A_i_*, *B_i_* ∈ *A_i_* and *A_j_*, *B_j_* ∈ *A_j_* is given by

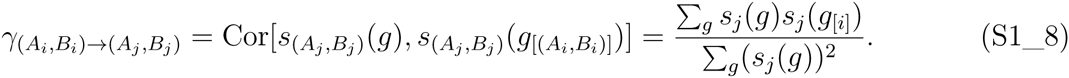

where 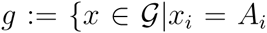 or *x_i_ = B_i_* and *x_j_* = *A_j_* or 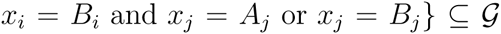 such that the sum is only calculated over the subset of genotypes carrying one of the two focal alleles at each focal locus. Thus, 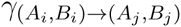 is a quadratic matrix of dimension 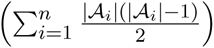. Note that in the case where 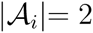 for all loci *i*, we obtain equation 9 in [14].

Likewise, the epistatic effect of a mutation in locus *i* with alleles (*A*_*i*_,*B*_*i*_) on other loci (and pairs of alleles) can be calculated as

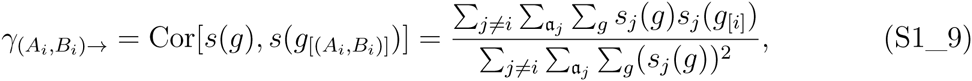

where the summation index 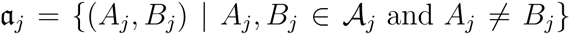 is over the set of subsets of size two that can be constructed from all alleles found at locus *j*. Note that the third summation index *g* changes depending on **a**_*j*_.

An additional summation allows calculation of the epistatic effect of a mutation in locus *i* carrying allele (*A*_*i*_) on other loci (and pairs of alleles) can be calculated as

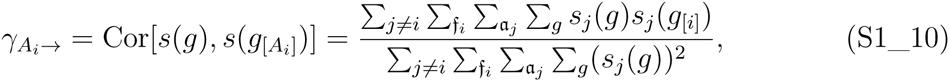

where 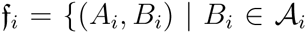 and *A_i_* ≠ *B_i_*} such that the sum is only calculated over the elements of the set of subsets of size two that can be constructed from all alleles found at locus *i* that contain allele *A*_*i*_.

Then, summing over 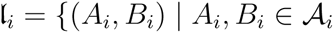 and *A_i_* ≠ *B*_*i*_}, i.e., the elements of the set of subsets of size two that can be constructed from all alleles found at locus *i*, gives the epistatic effect of a mutation in locus *i*

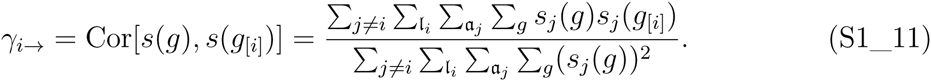

Similarly, the epistatic effect of other mutations (again considering pairs of alleles first) on locus *j* with alleles (*A*_*i*_, *B*_*i*_) can be calculated as

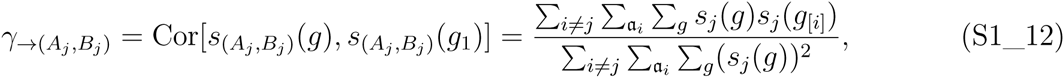

the epistatic effect of other mutations on locus *j* carrying allele (*A_j_*) is given by

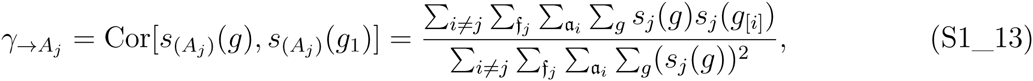

and the epistatic effect of other mutations on locus *j* becomes

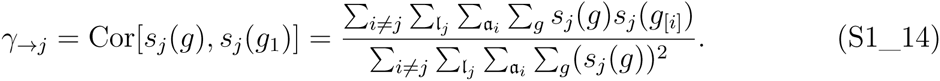

Finally, *γ_d_*, that is the decay of correlation of fitness effects with Hamming distance *d*, (i.e., the cumulative epistatic effect of *d* mutations averaged over the entire fitness landscape) is calculated as

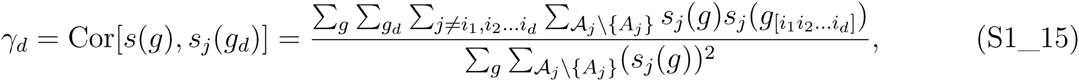

where the last summation is over all different alleles present at locus *j* except the one carried by genotype *g* at locus *j*. Note that there is unfortunately no multi-allelic analog to equation (14) of [14]. Furthermore, as desired when there are only two alleles at a given locus, equations (S1_9 – S1_11) and equations (S1_12 – S1_14) collapse and give identical values, and reduce to their di-allelic counterparts (i.e., eq. 7-8 in 14).

### Fraction of epistasis

The second statistic quantifies whether specific pairs of alleles between two loci interact epistatically, and if so whether they display magnitude epistasis (i.e., fitness effects are non-additive, but fitness increases with the number of mutations), sign epistasis (i.e., one of the two mutations considered has an opposite effect in both backgrounds) or reciprocal sign epistasis (i.e., if both mutations show sign epistasis; [34, 35]; for an implementation see [36]).

Using equation (S1_1) the type of epistatic interaction between a mutations at locus *i* and *j* (with *i* ≠ *j*) when introduced on some reference genotype 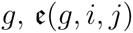, can formally be given as

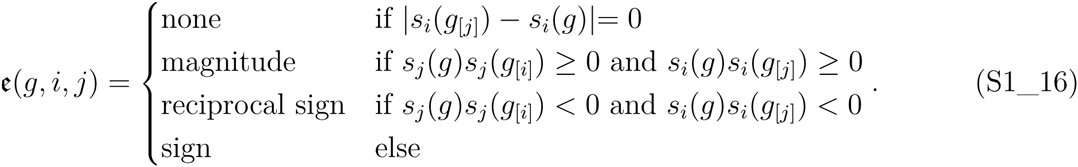

Note that for numerical purposes we allowed for a very small deviation *ε =* 10^−6^ in the case of no epistatic interaction. For ease of notation, we treat the elements in 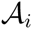 as fixed and ordered, and define **I**(*g*_[*i*]_) as the index of the allele present at locus *i* in genotype *g* in 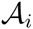 such that the fraction of epistasis over all 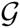 can be calculated as

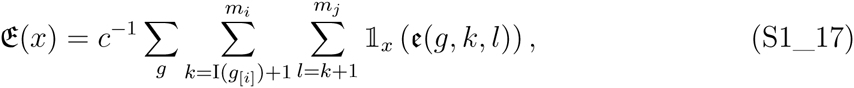

where *x* ∈ {none, magnitude, sign, reciprocal sign}, 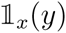 is the indicator function that is 1 if *x* = *y* and 0 otherwise, and 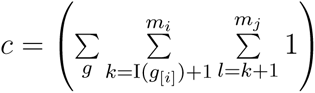 is a normalization constant.

## Supporting Information 2: Overview of different fitness landscape models introduced in the main text

In the following, we briefly introduce the various fitness landscape models introduced and discussed in the main text and provide references to selected relevant literature. For an excellent general overview, we refer the reader to a review by [10]

### The additive model

In the additive model the log-fitness of each genotype is given by the sum over the individual per locus log-fitness effects. Thus, the fitness effect of a specific allele – drawn from a Normal distribution with mean *μ*_*a*_ and variance 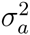 [14] [see also 54] – is independent of its genetic background (i.e., it is constant across all genetic backgrounds), such that all mutations are non-interacting (i.e., there is no epistasis) and the resulting single-peaked fitness landscape is (maximally) smooth. In particular, the roughness-to-slope ratio is 0 and 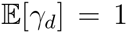 for the entire range of mutational distances *d*. Note that when fitnesses are not measured on log-scale, the fitness of a genotype is simply the product over the individual fitness effects, which is why this model is sometimes also referred to as the multiplicative model.

### The House-of-cards model

On the other extreme, in the House-of-cards (HoC) model [53] the fitness of each genotype is an i.i.d. normally distributed random variable with zero mean and variance 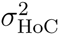 resulting in an uncorrelated, maximally rugged fitness landscape that is characterized by multiple fitness maxima and minima [55, 56]. In particular, the fitness effect of an allele entirely depends on its genetic background such that there is complete interaction between all loci (i.e., full epistasis) which is also reflected in the roughness-to-slope ratio and 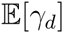 that become infinity and zero (for *d* > 0), respectively.

### The Rough Mount Fuji model

Introduced by [57], the Rough-Mount-Fuji (RMF) model – named after the eponymous mountain in Japan –, which was initially formulated in the context of protein evolution [see also 54, for a simplified version], interpolates between the former two extremes. The fitness of a genotype is composed by an additive component (parametrized by *μ*_*a*_ and 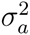; see above) and a HoC component (parametrized by 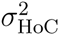) such that the extent of epistatic interactions ranges between none (additive model) to complete (HoC model) depending on the relative sizes of these three parameters. In particular, when 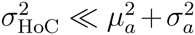 the RMF model becomes an additive model whereas for 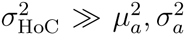 it essentially behaves like a HoC model [14]. Accordingly, 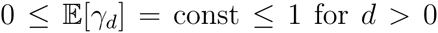 (see Figure 4a) and the roughness-to-slope ratio ranges from zero to infinity.

### The Kauffman NK model

Another frequently used fitness landscape model that, as the RMF model, also interpolates between the additive and the HoC model [55, 56] is the Kauffman NK model, where *N* di-allelic loci interact with *K* ∈ {0,1, …, *L* − 1} randomly assigned other loci. In particular, for *K* = 0 the NK model collapses to an additive model whereas for *K* = *L* − 1 it approaches the HoC model. Although there are different ways how groups of interacting loci can be chosen [58], properties such as the mean number and height of local optima tend only to be weakly dependent on the exact choice being made. For the NK model, 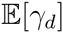 is a non-negative monotonically decreasing function in *d*, and the roughness-to-slope ratio can again range from zero to infinity.

### The eggbox model

Introduced by [14], the eggbox model is a maximally epistatic, anticorrelated fitness landscape model (i.e., all loci interact with each other up to interactions of order *L*), in which the fitness effect of an allele switches from the highest to lowest value (or vice versa) between genetic backgrounds one step apart. Accordingly, depending on whether two genotypes are separated by an odd or even Hamming distance, their absolute fitness difference is either twice the mean allelic fitness effect or zero. Thus, this model generates an extreme case of reciprocal sign epistasis in which each mutation is either deleterious or compensatory, multiple local optima exist, and *γ*_*d*_ accordingly oscillates between −1 and 1.

## Supporting Information 3: Supporting Figures

### Supporting Information 4: Addressing measurement uncertainty in fitness landscape statistics

**Figure S3_1.**
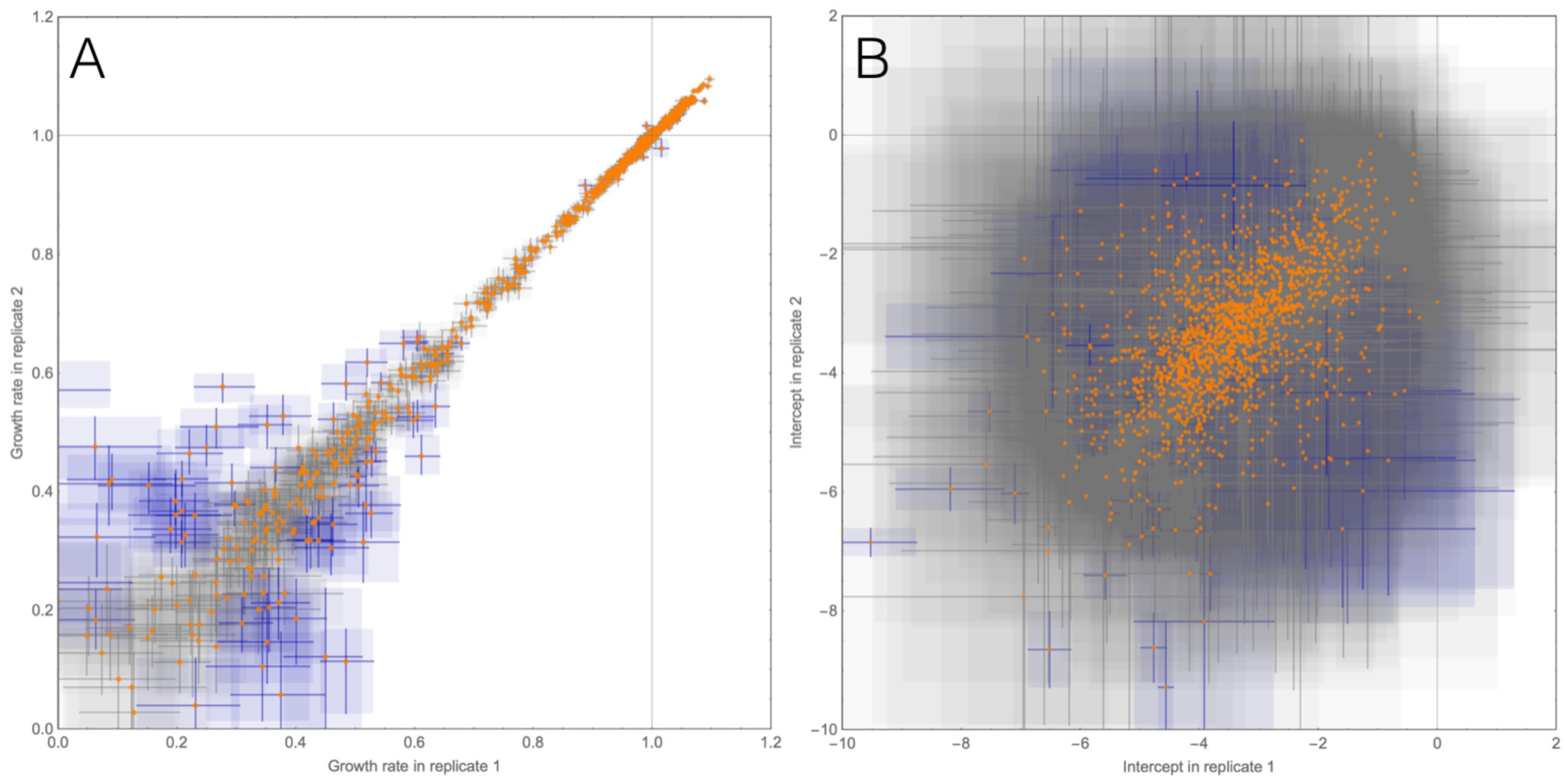
(A) Scatterplot demonstrating the correlation of estimated growth rates (*R*^2^ = 0.944). Orange dots indicate median growth rates, rectangles indicate the limits of the 95% credibility intervals (in blue if they do not overlap between replicates). (B) In contrast, estimated initial population sizes are much more variable between replicates, demonstrating that initial representation in the mutant library had no effect on the estimated growth rate.

**Figure S3_2.**
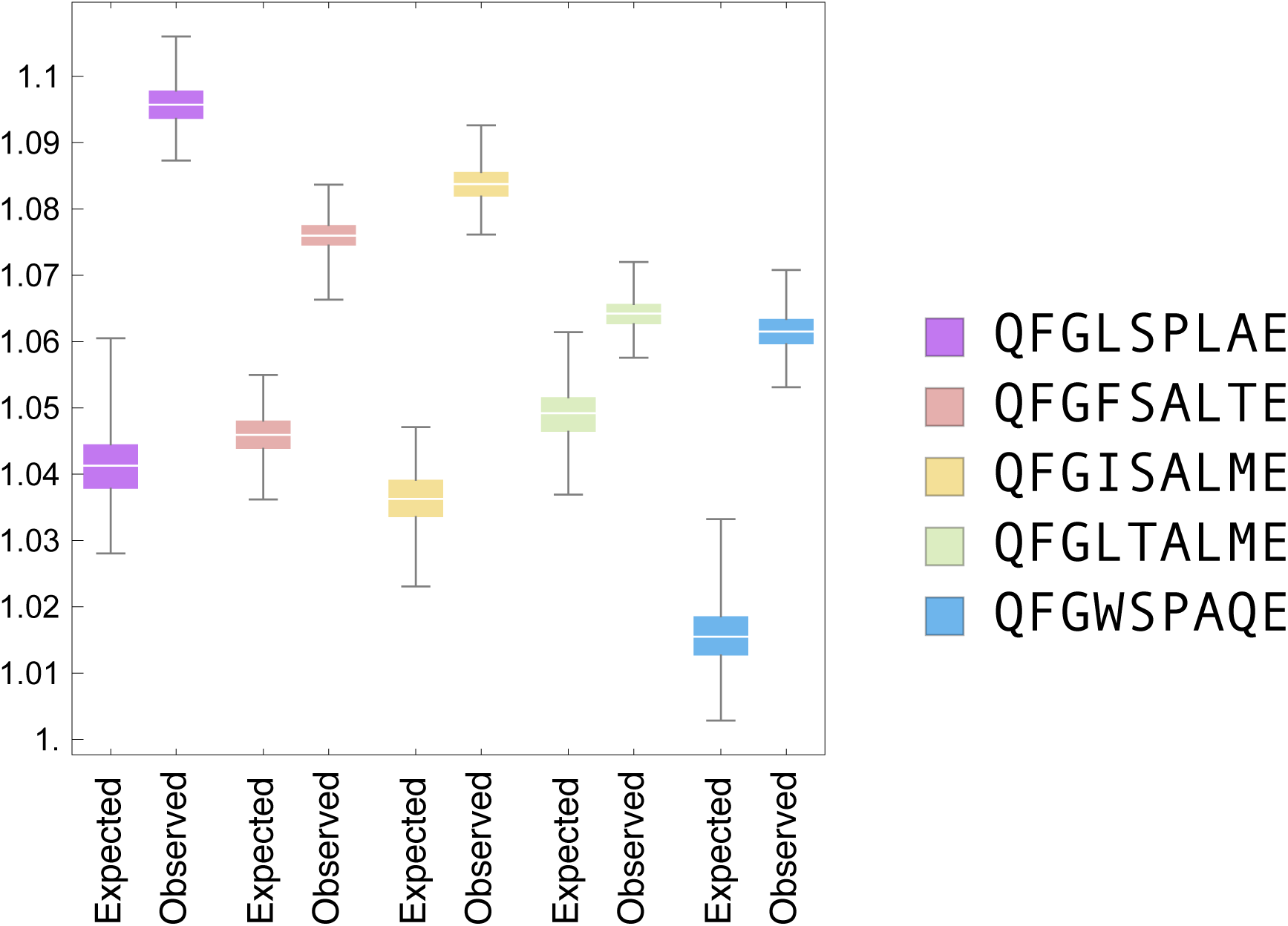
Expected versus observed fitness for all 5 beneficial local optima of the fitness landscape, obtained from 1000 posterior samples.

**Figure S3_3.**
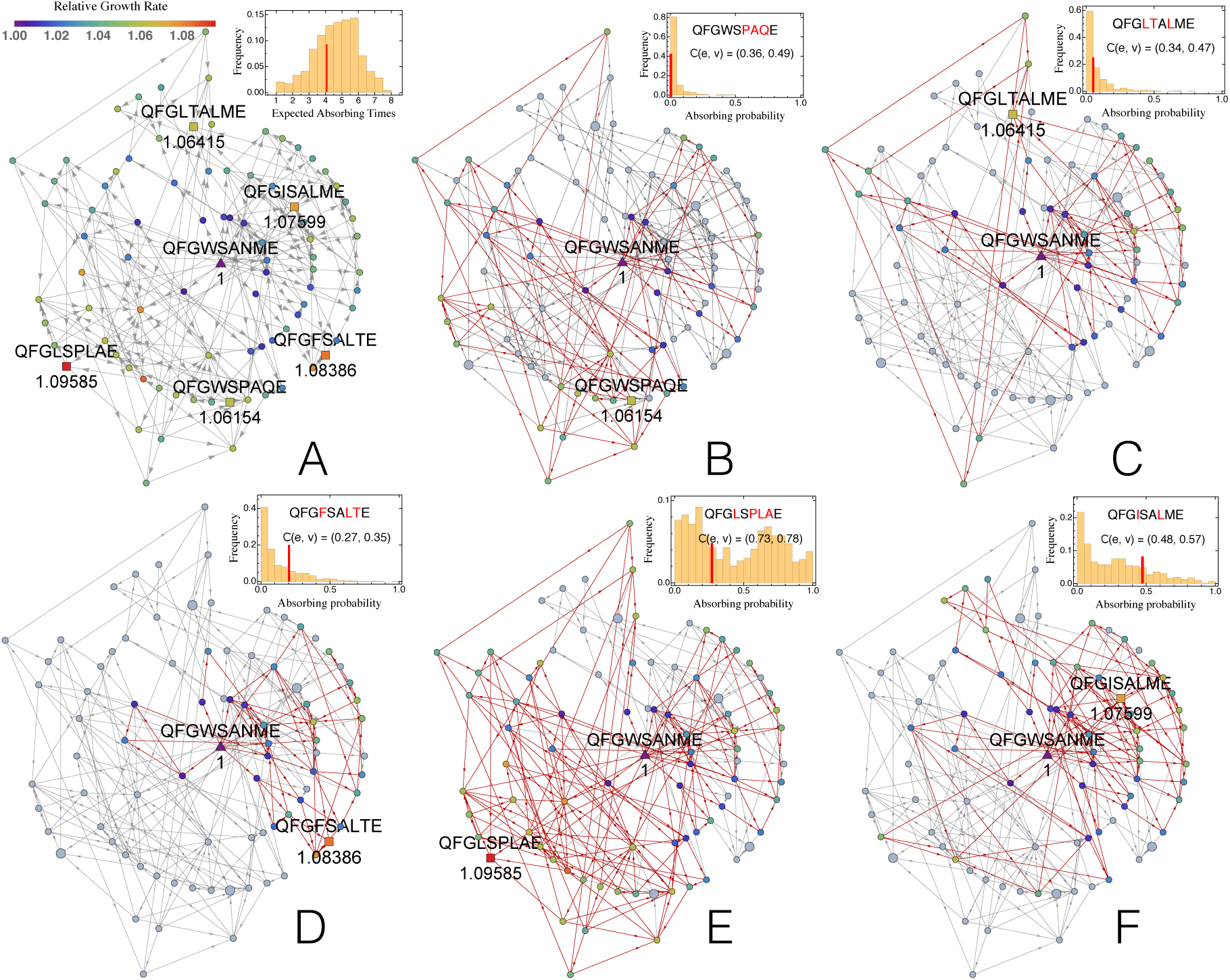
Graphical illustration of the sub-landscape of all beneficial mutants with respect to the parental type. (A)Each vertex (circle) corresponds to a mutant that is beneficial with respect to the parental type (triangle), colored according to their fitness. Arrows connect mutants that differ by a single substitution, with their direction indicating an increase in fitness. The five local optima in this sub-landscape are indicated as squares. Inset: distribution of average lengths of adaptive walks starting from any type in the full landscape (i.e., absorbing times of the Markov chain). The red line indicates mean path lengths for adaptive walks from the parental genotype. (B)-(F) Paths leading towards the different optima – highlighted by their sequence name – when beginning an adaptive walk at the parental genotype. Inset (B) - (F): Distribution of absorbing probabilities, that is, the probability to reach a specific optimum starting from a given mutant computed for all mutants in the data set. The red line corresponds to the respective probability when starting from the parental sequence. The global optimum (panel E) is in general reached with a very high probability, but there are starting points from which it is poorly accessible. *C*(*e*,*v*) quantifies the landscape coverage: although 73% of all edges and 78% of all vertices can be included in walks from the parental genotype to the global optimum, it is reached in only 26% of all possible walks. Conversely, 47% of walks will reach the local optimum in panel C, which is lower in fitness, but closer to the parental type in terms of its Hamming distance. This demonstrates that evolution on this landscape is not well predicted by fitness only, and that a population may become stuck in a sub-optimal fitness peak.

**Figure S3_4.**
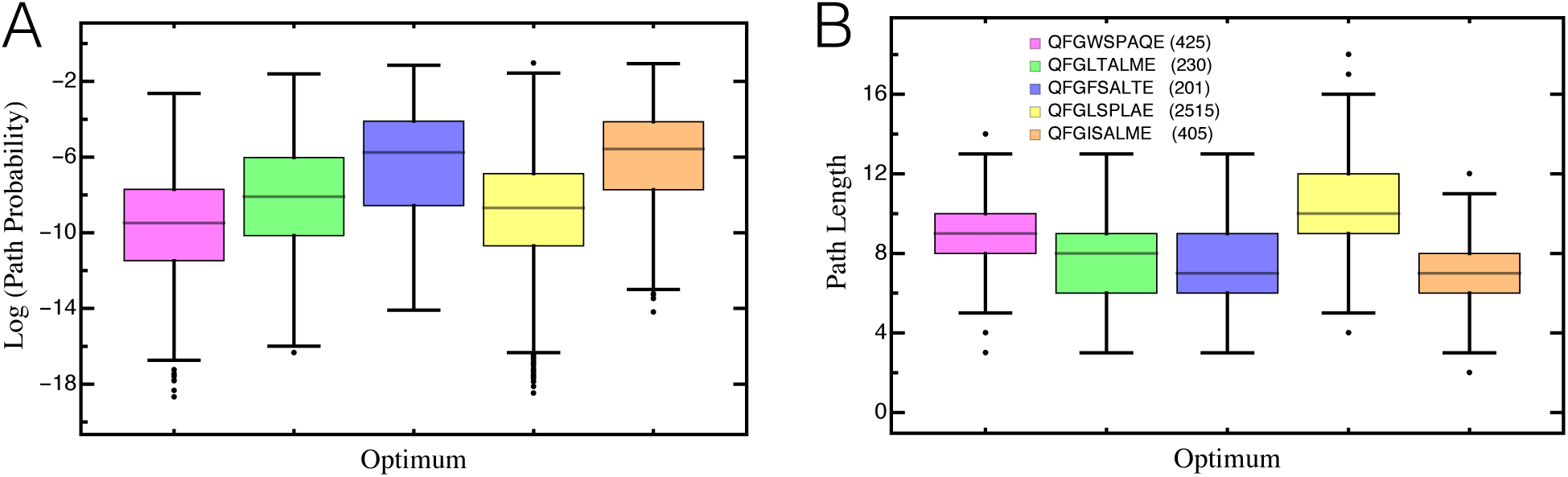
(A) Path probability of reaching a specific local optimum when beginning an adaptive walk at the parental genotype. Results are presented on log scale. Boxes represent the interquartile range (i.e., the 50% C.I.), whiskers extend to the highest/lowest data point within the box ± 1.5 times the interquartile range, and black circles represent close outliers. (B) Path length (i.e., the number of substitutions) to reach a specific local optimum. Inset: Color coding for the different optima with the total number of different paths leading to the respective optimum in brackets. There is a total of 3776 different paths that can be taken from the parental genotype, 66% of which lead to the global optimum (yellow). Most of these, however, require multiple substitutions, such that the probability to reach the global optimum is generally low. In contrast, the local optimum close to the parental genotype (orange) can be reached with only a few substitutions, and thus, with relatively high probability.

**Figure S3_5.**
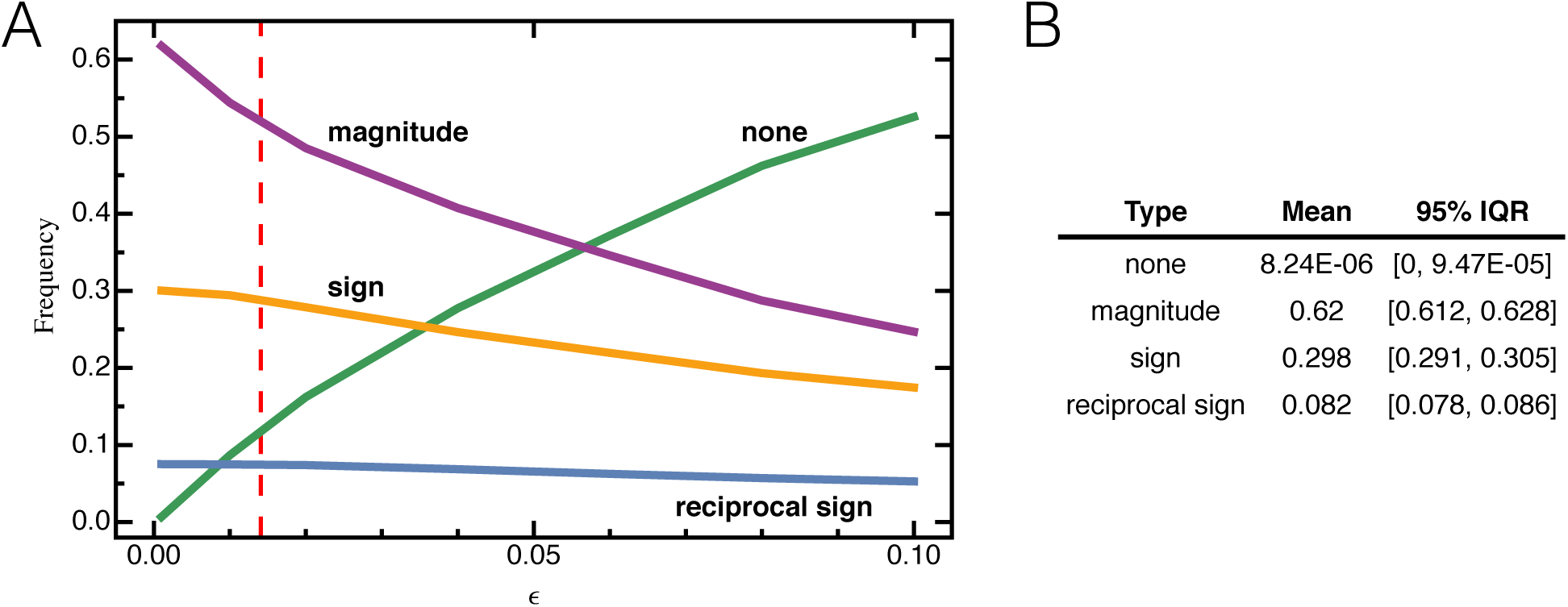
(A) The fraction of epistasis (eq. S1_17) calculated for different values of *ε*, quantifying the error threshold beyond which interactions between different alleles are considered to be epistatic. Expectedly, as *ε* increases the number of non-epistatic interactions between alleles increases, which in the majority of cases were previously classified as magnitude epistatic interactions. Likewise, the fraction of sign and reciprocal sign epistasis decreases. The red dashed line shows the mean absolute difference between median growth rates obtained from the two independent replicates calculated over all mutants with median growth rate larger than the mean median growth rate estimated for the substitution F583* (i.e., when substituting a stop codon) between the two replicates. Using this threshold for *ε*, we find that approximately 10% of all interactions are additive, approximately 50% show magnitude epistasis, whereas the sign and reciprocal sign epistasis remains roughly constant at 30% and 8%, respectively. (B) The mean and the 95% interquantile range (IQR) of the fraction of epistasis calculated over 10,000 posterior samples obtained from the MCMC simulations using the standard threshold *ε* = 10^−6^. There is only little to no variation in the fraction of epistasis and the mean value is essentially identical to the proportions reported in the main text, indicating robustness of the inferred fraction of epistasis, and confirming that variation in posterior samples is typically small (i.e., growth rates are estimated with high precision).

**Table S4_1.**
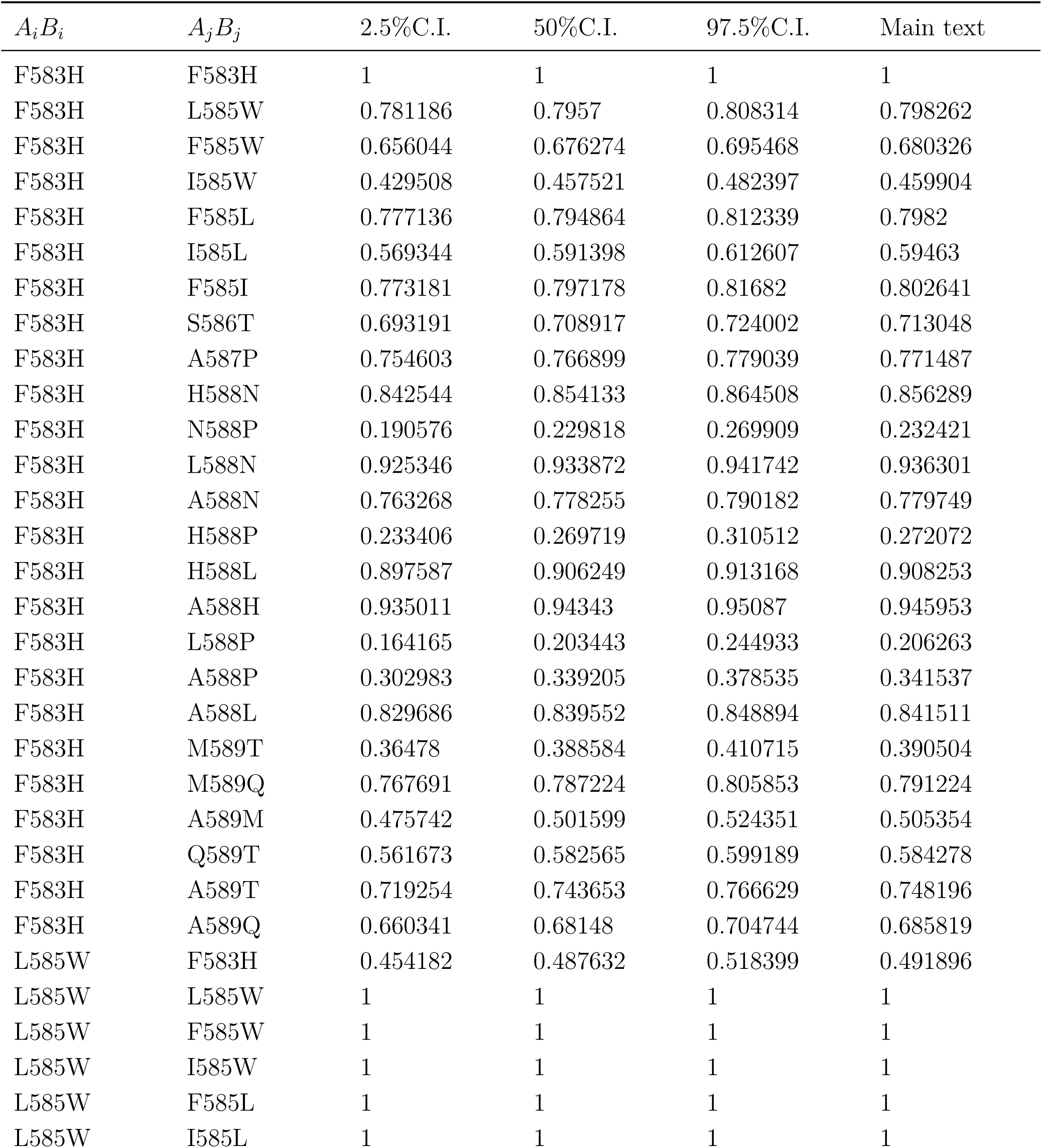
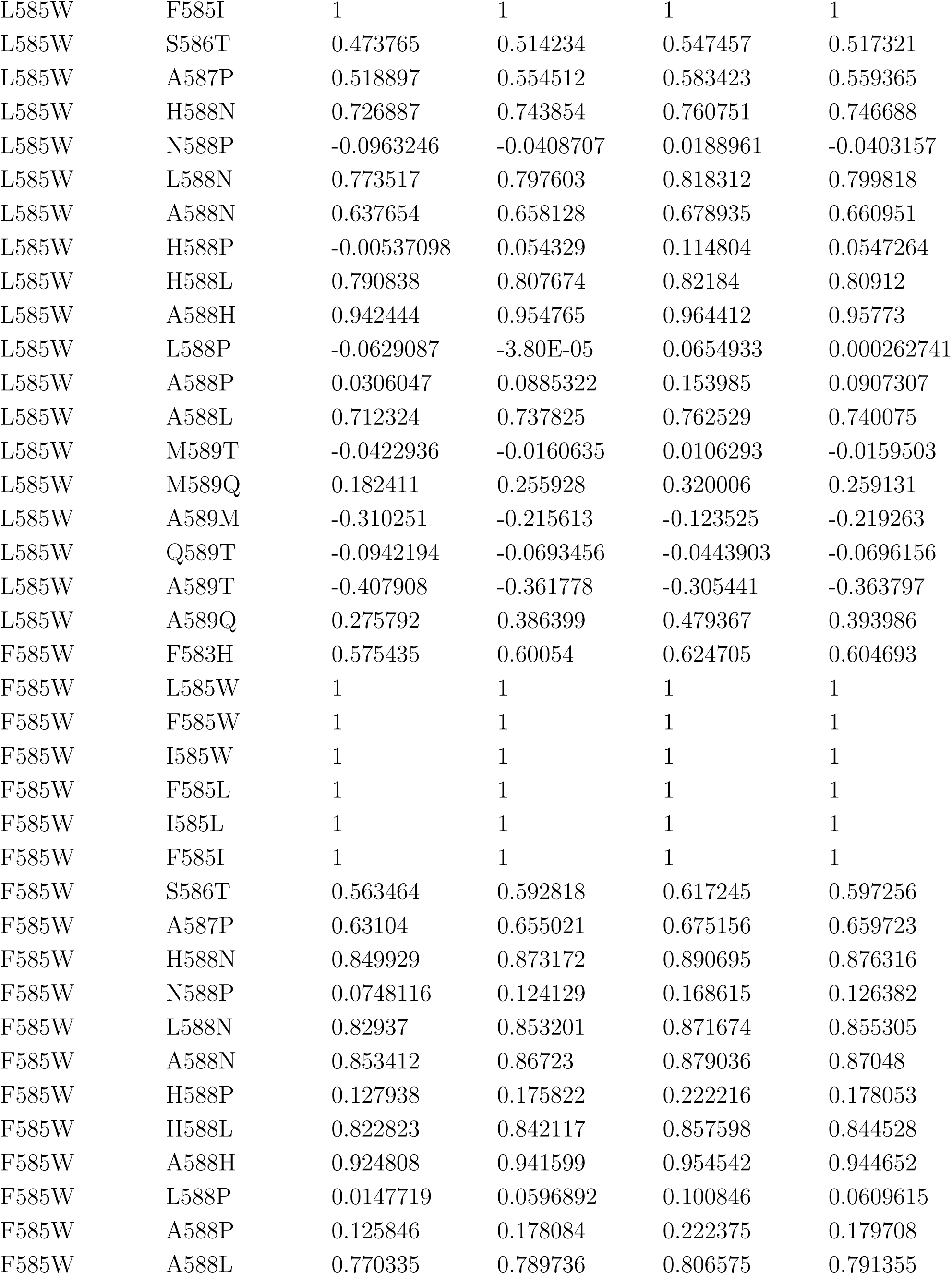
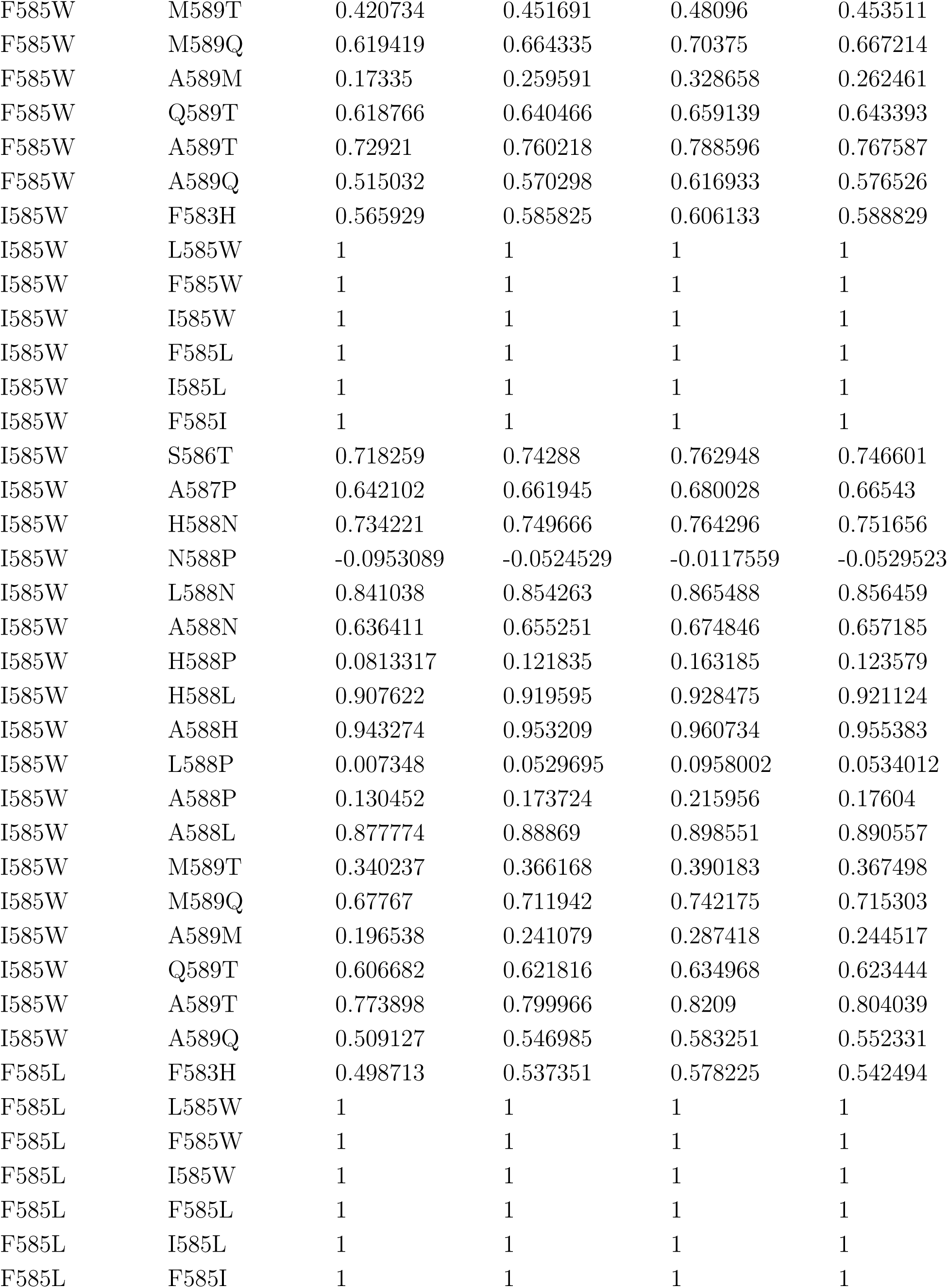
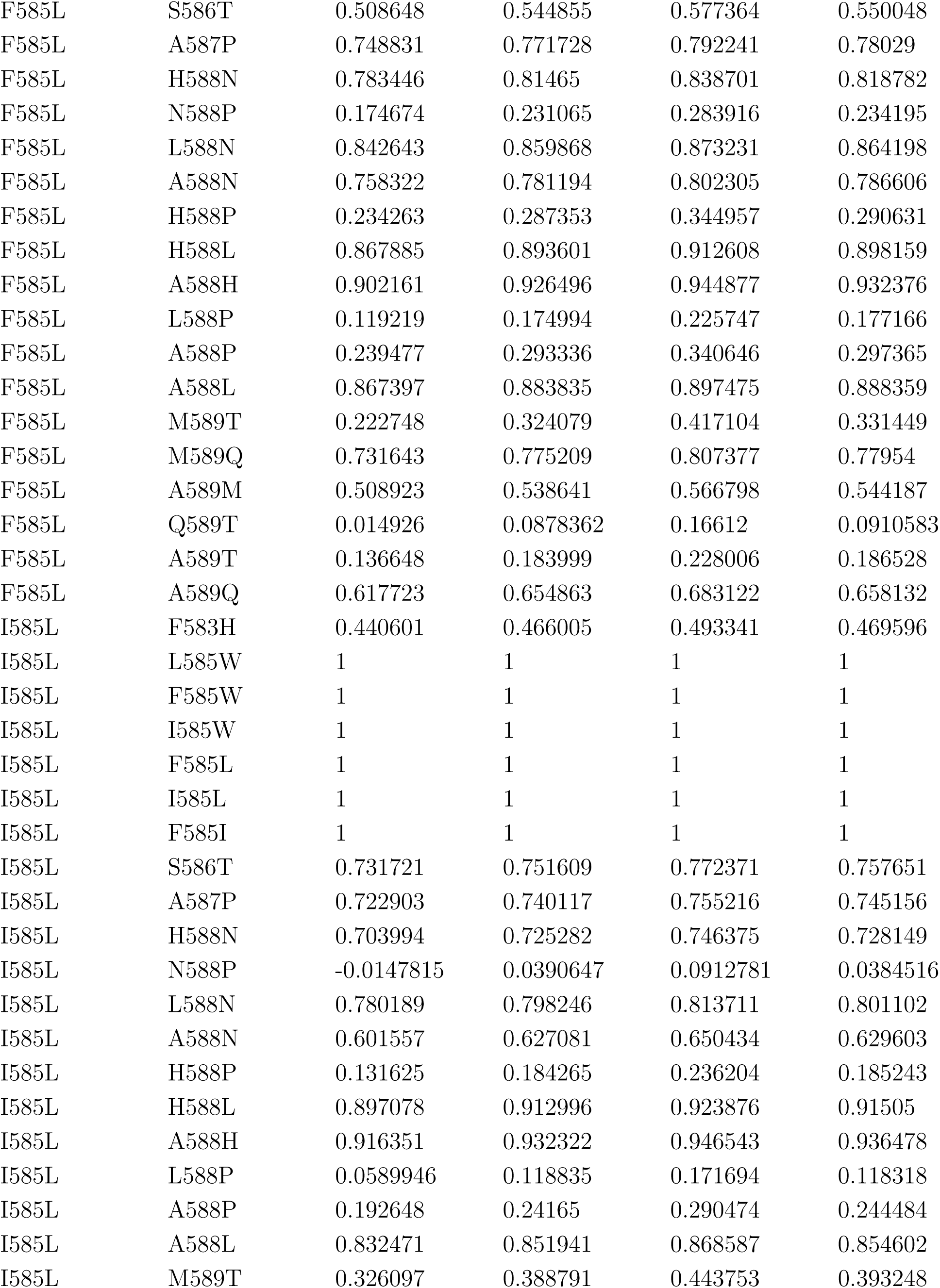
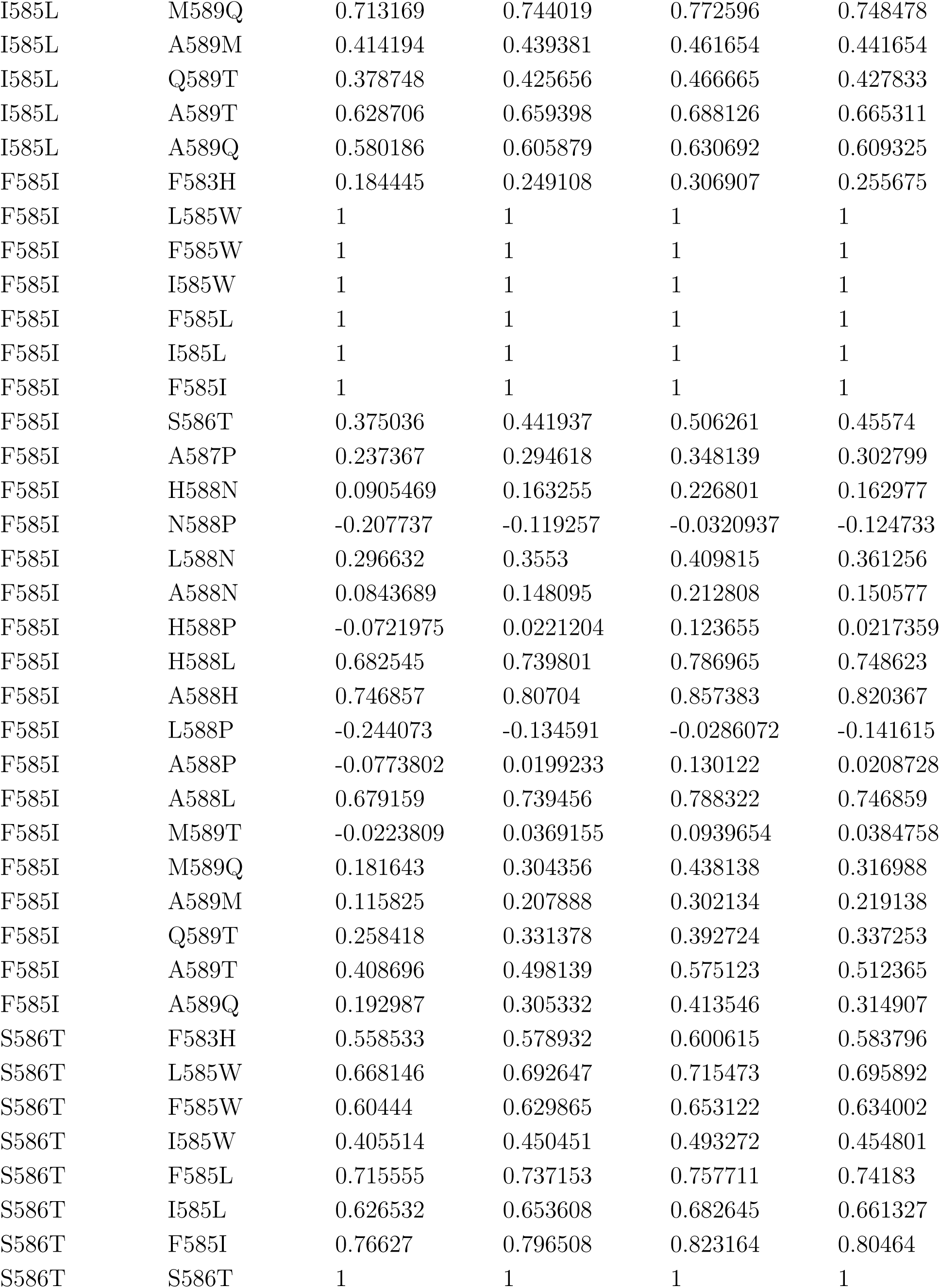
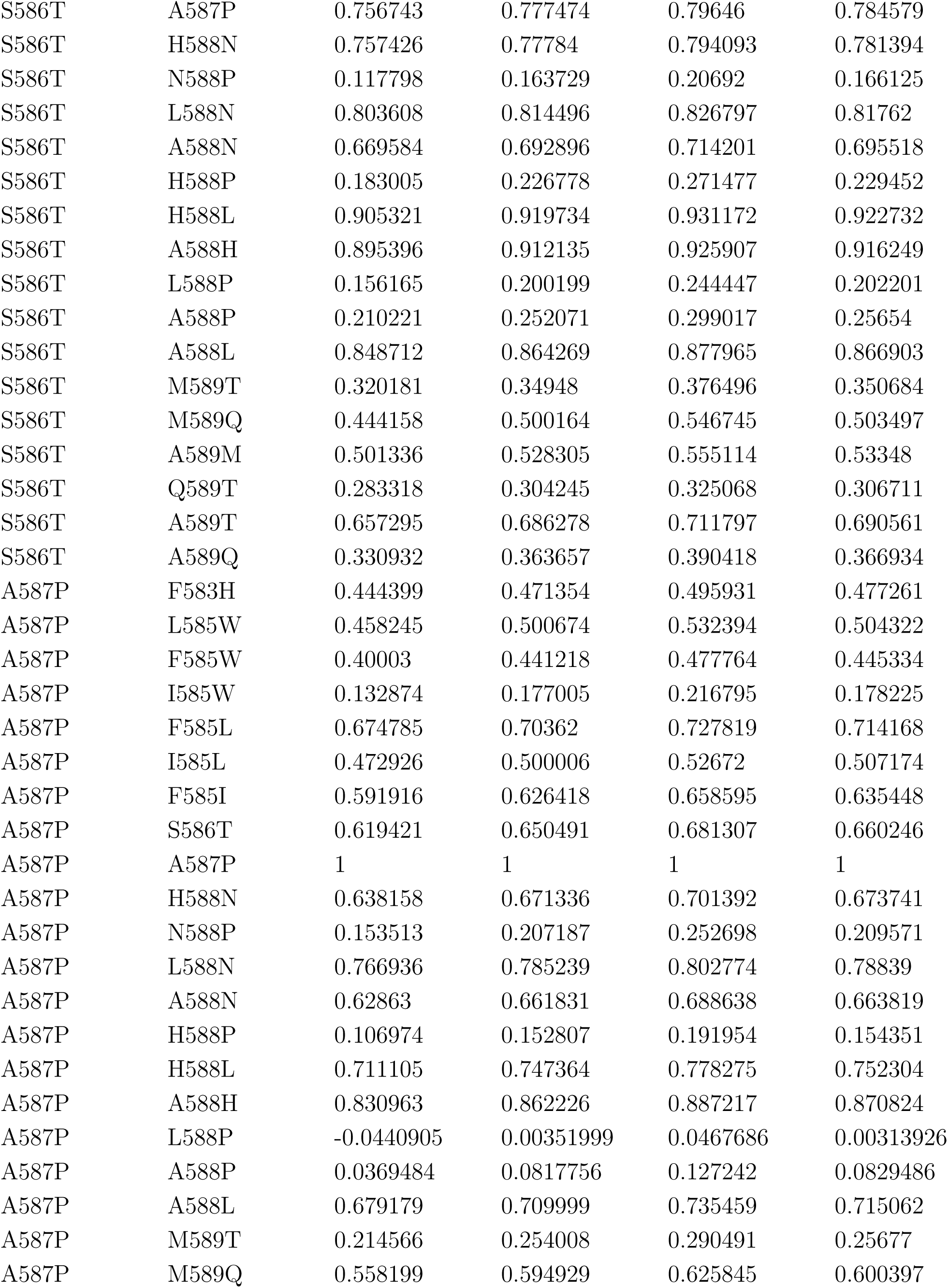
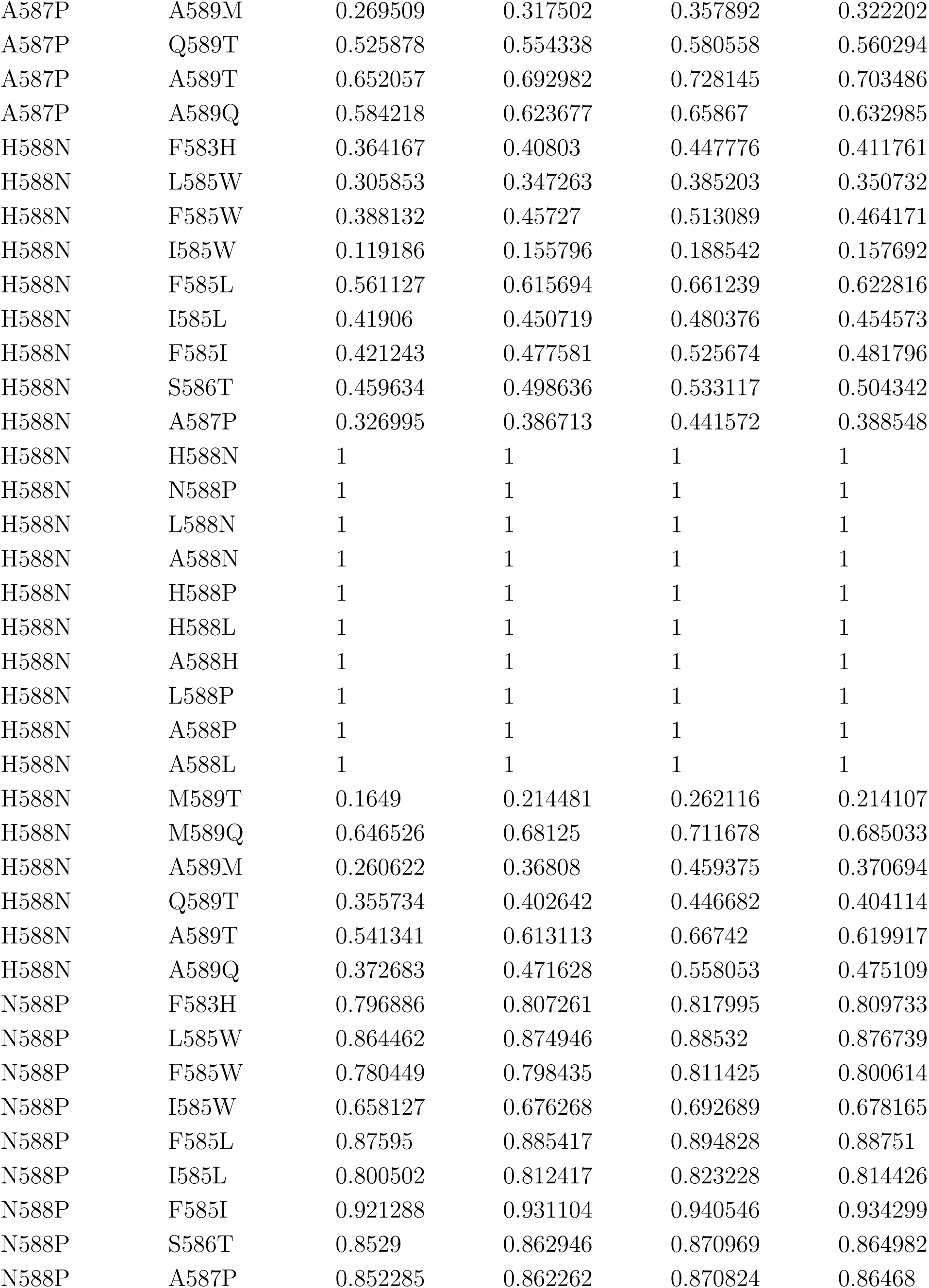
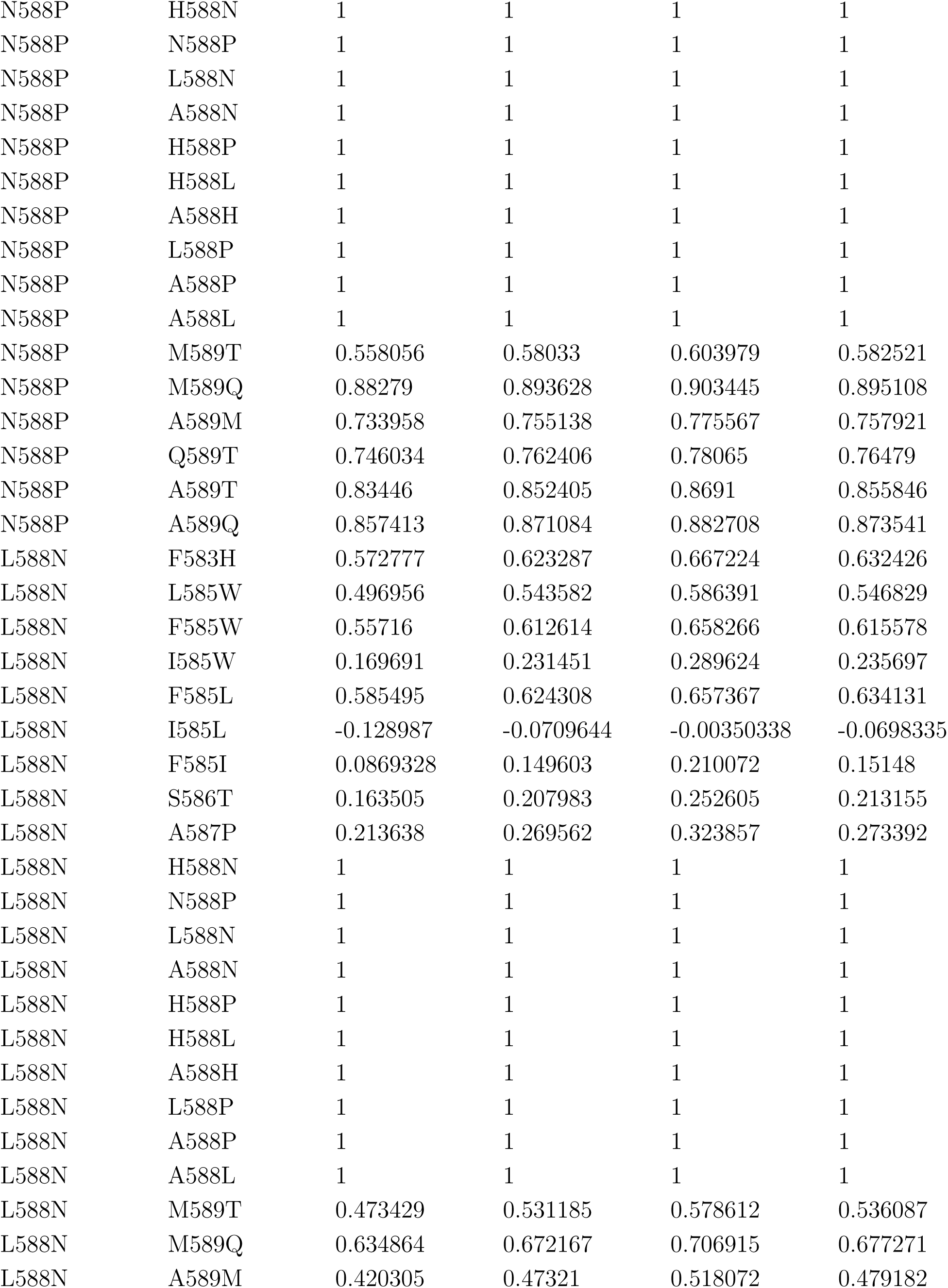
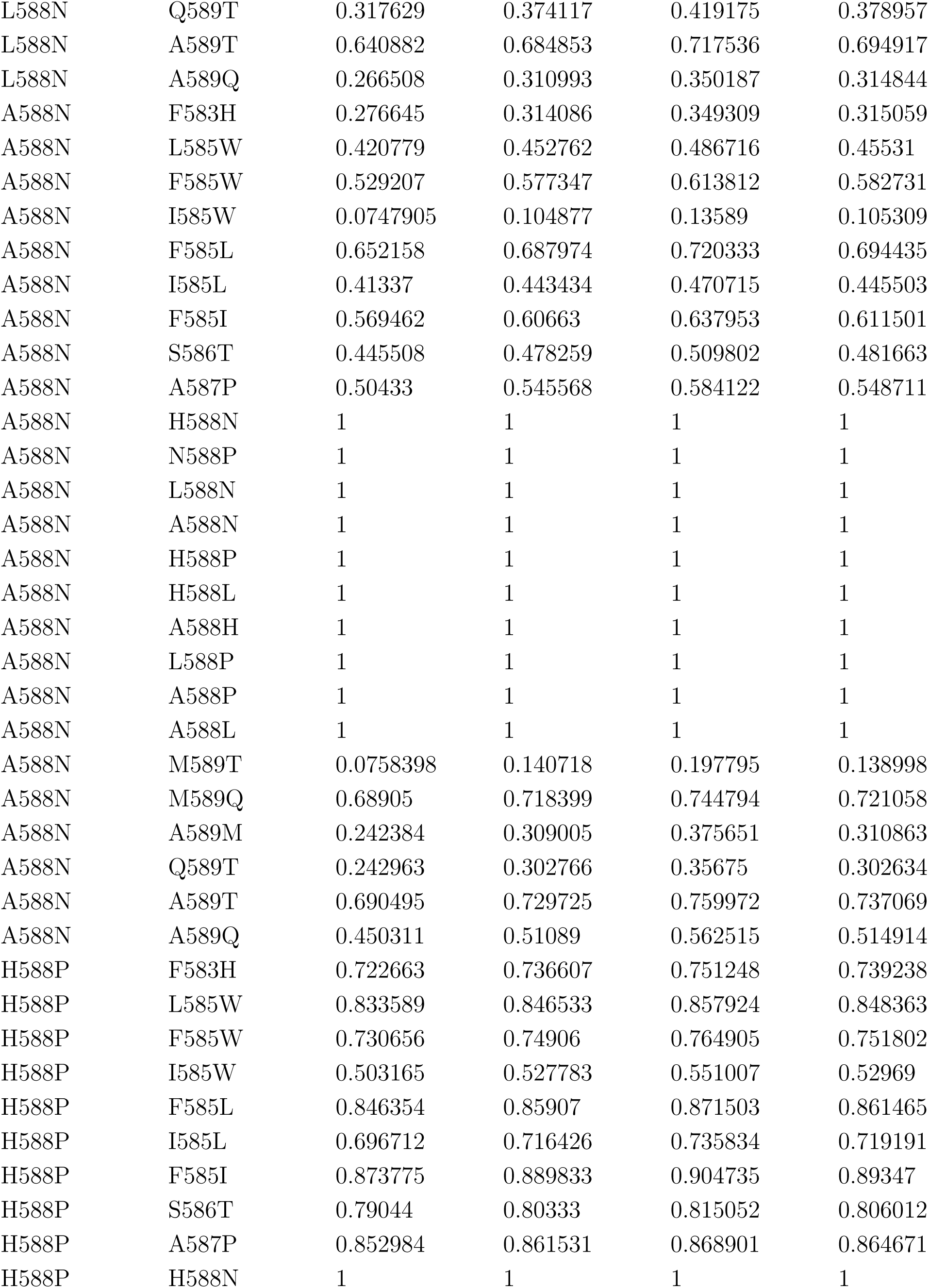
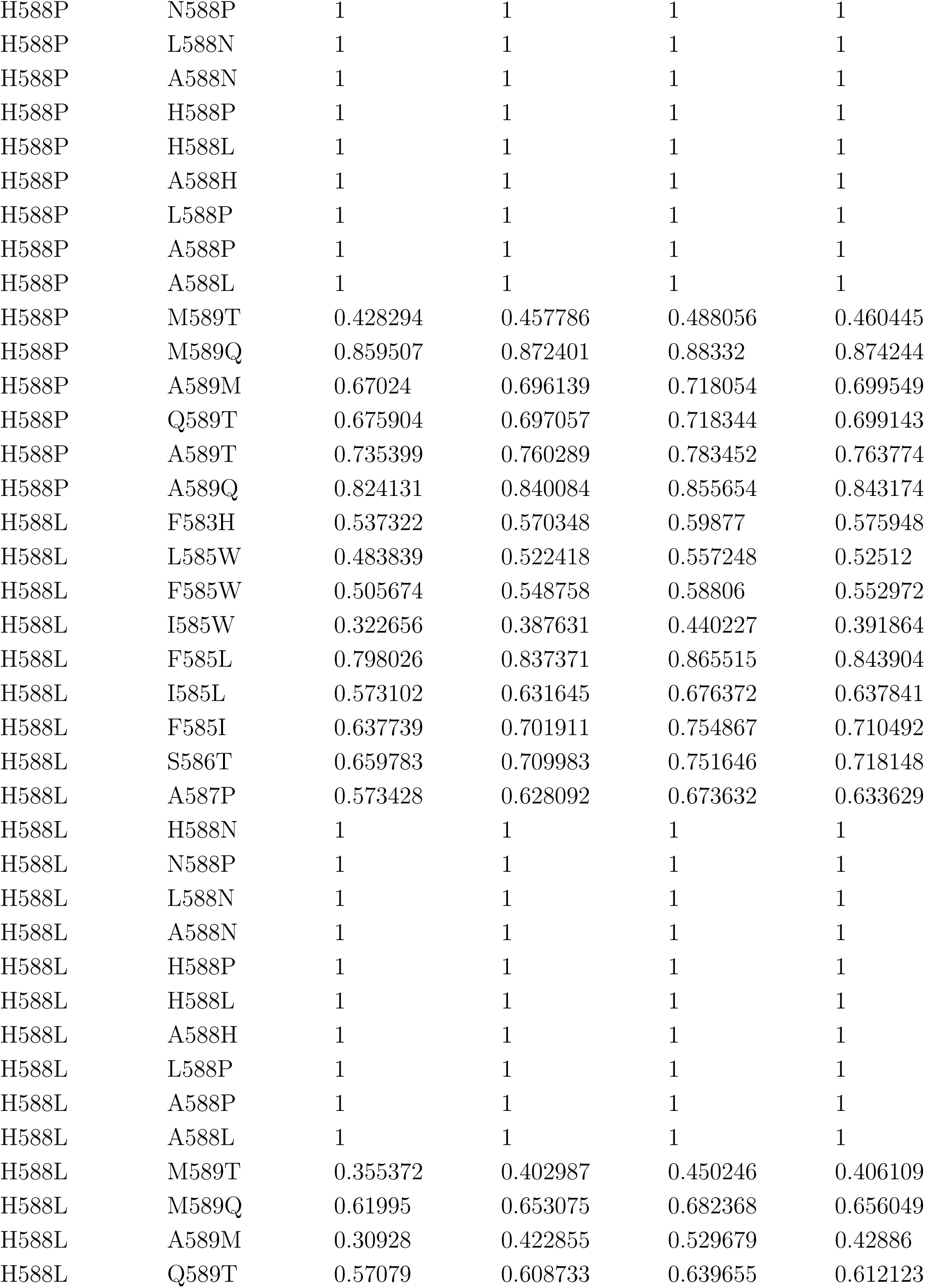
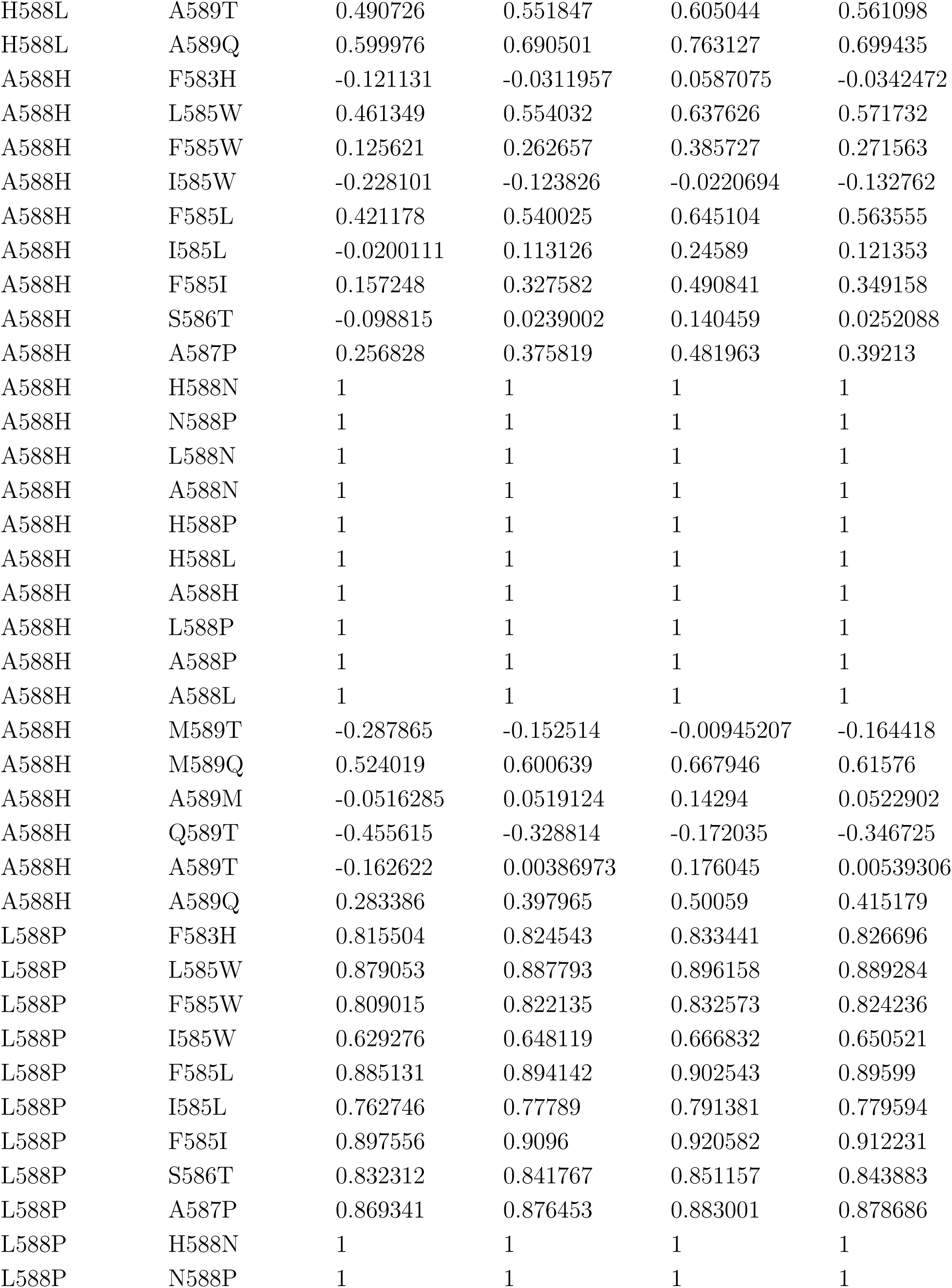
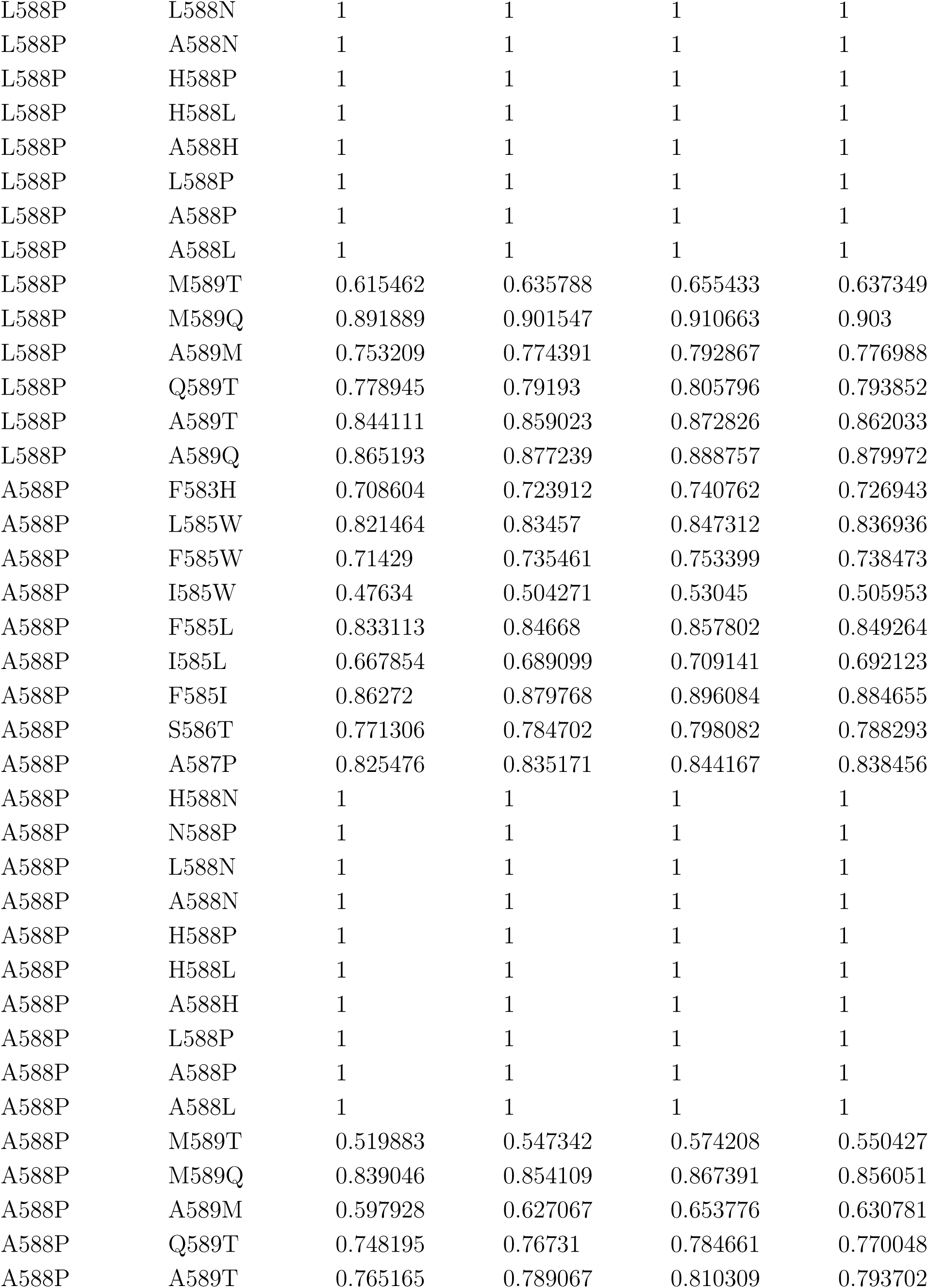
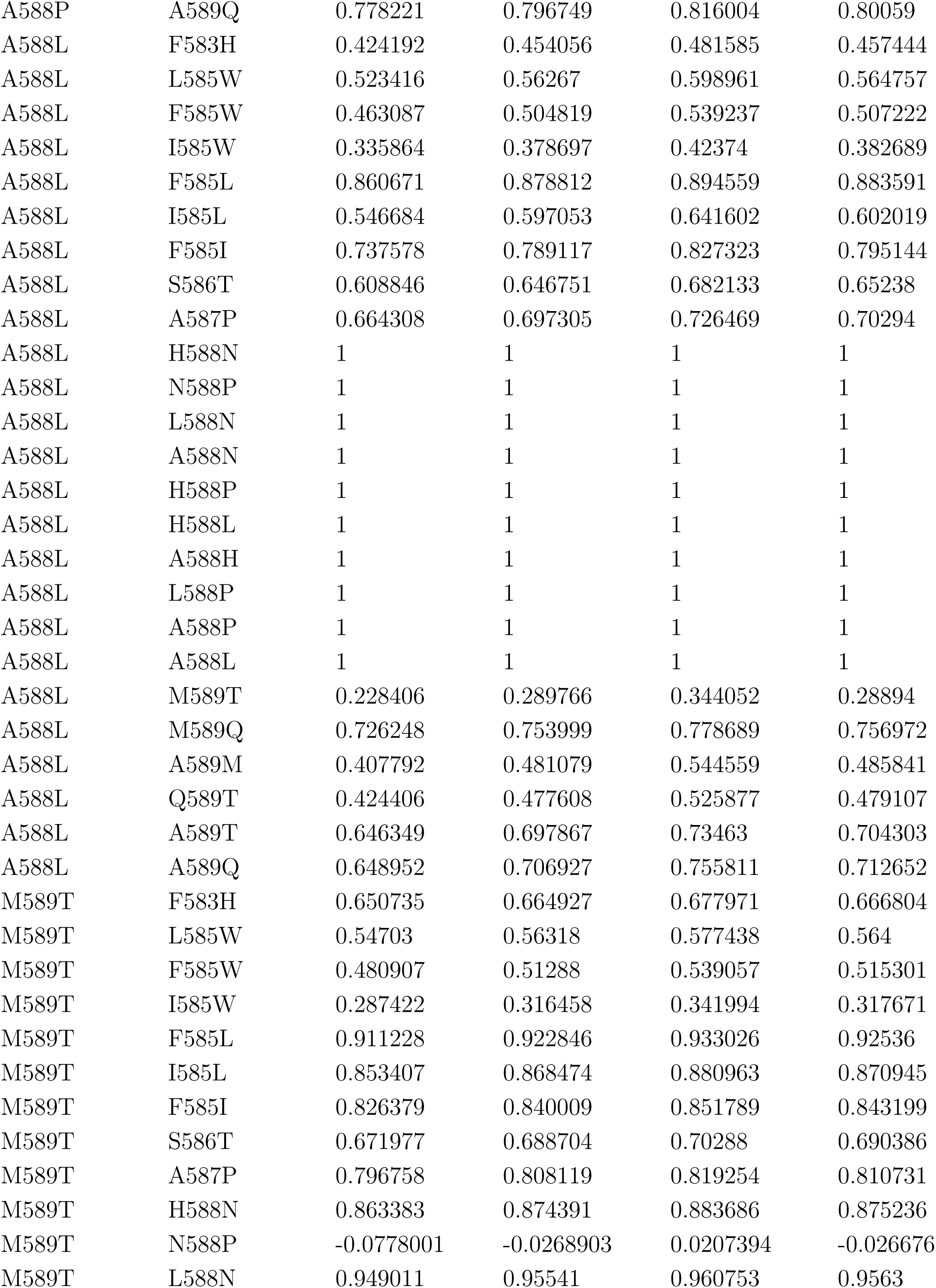
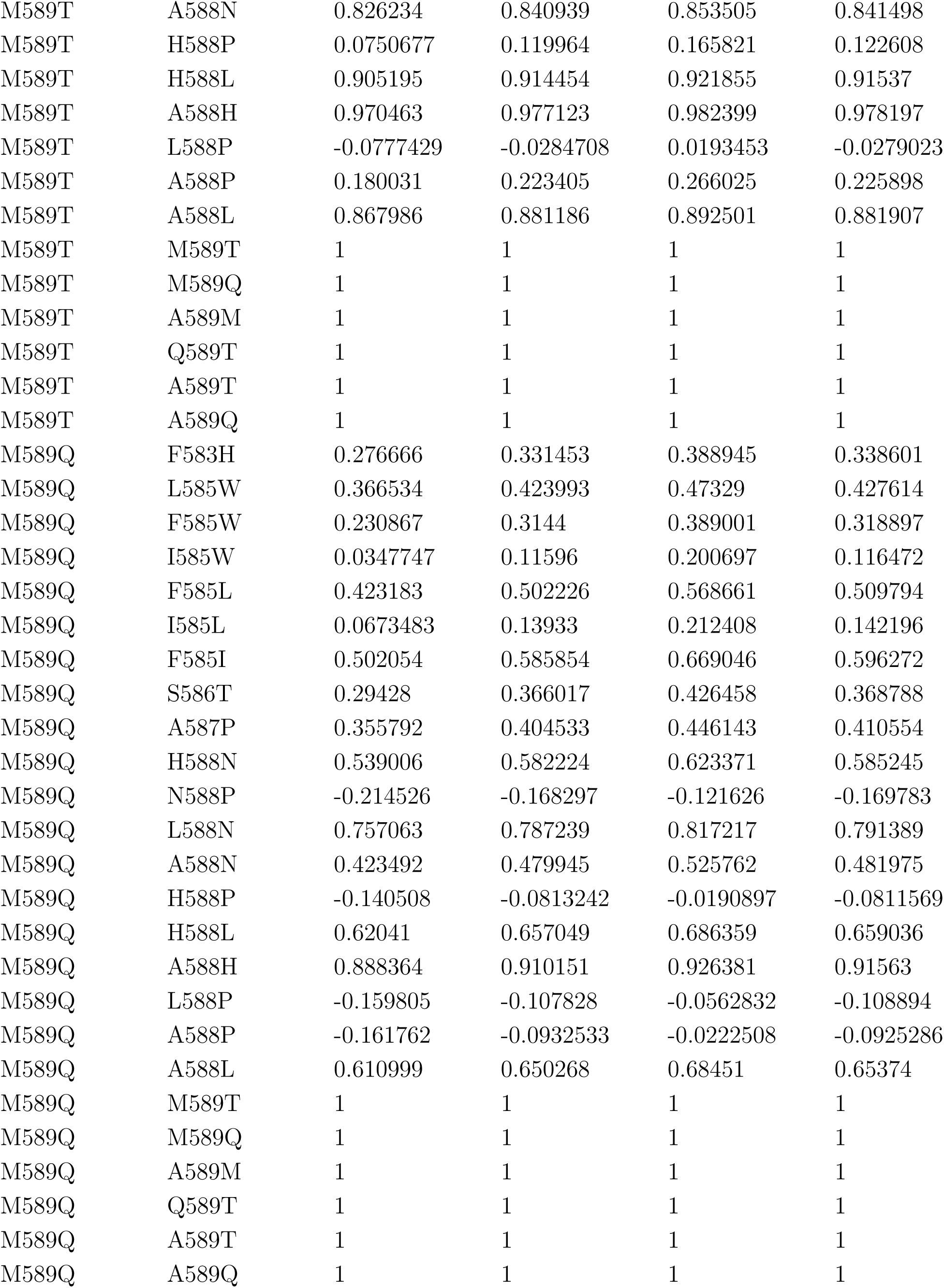
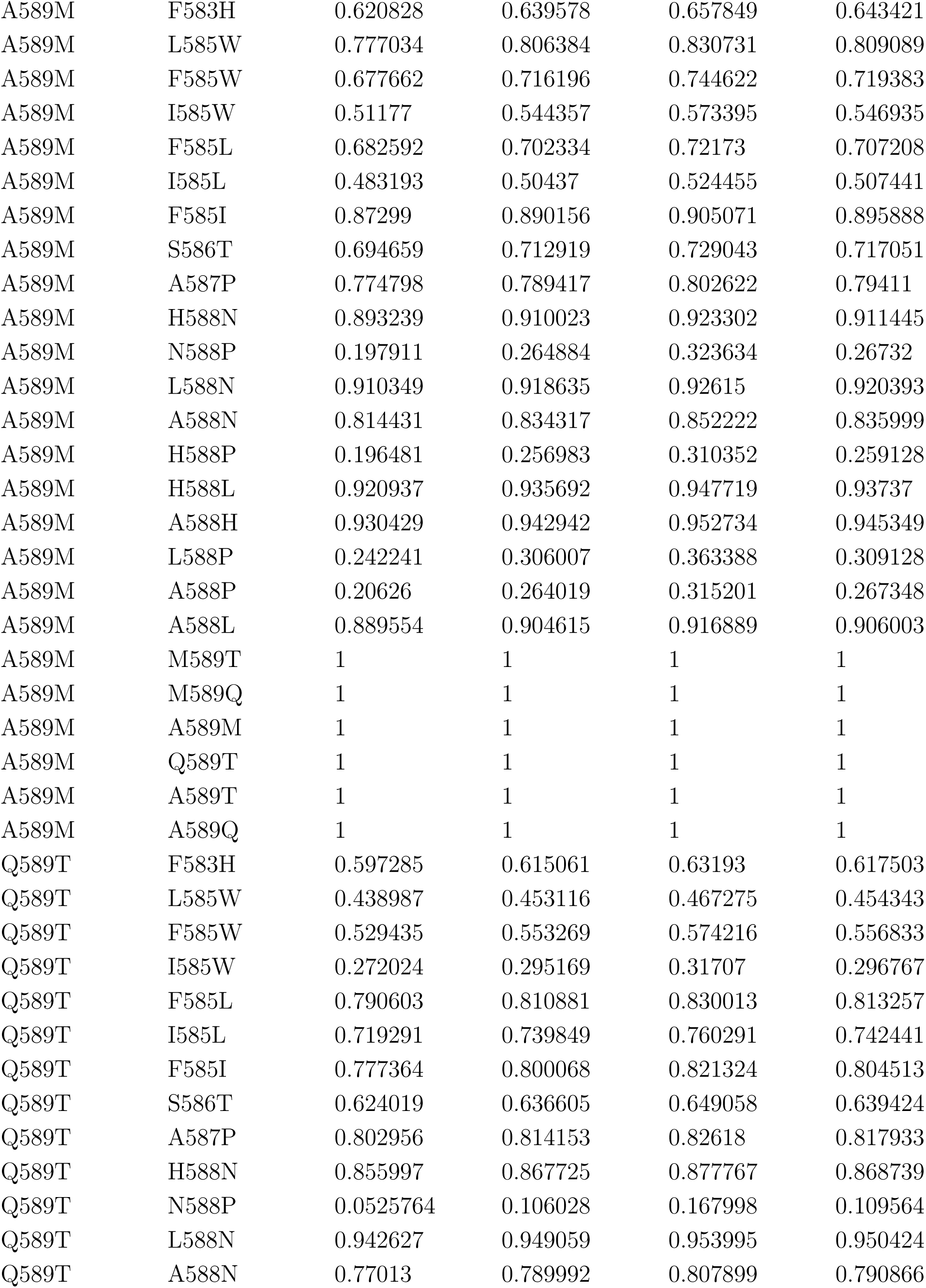
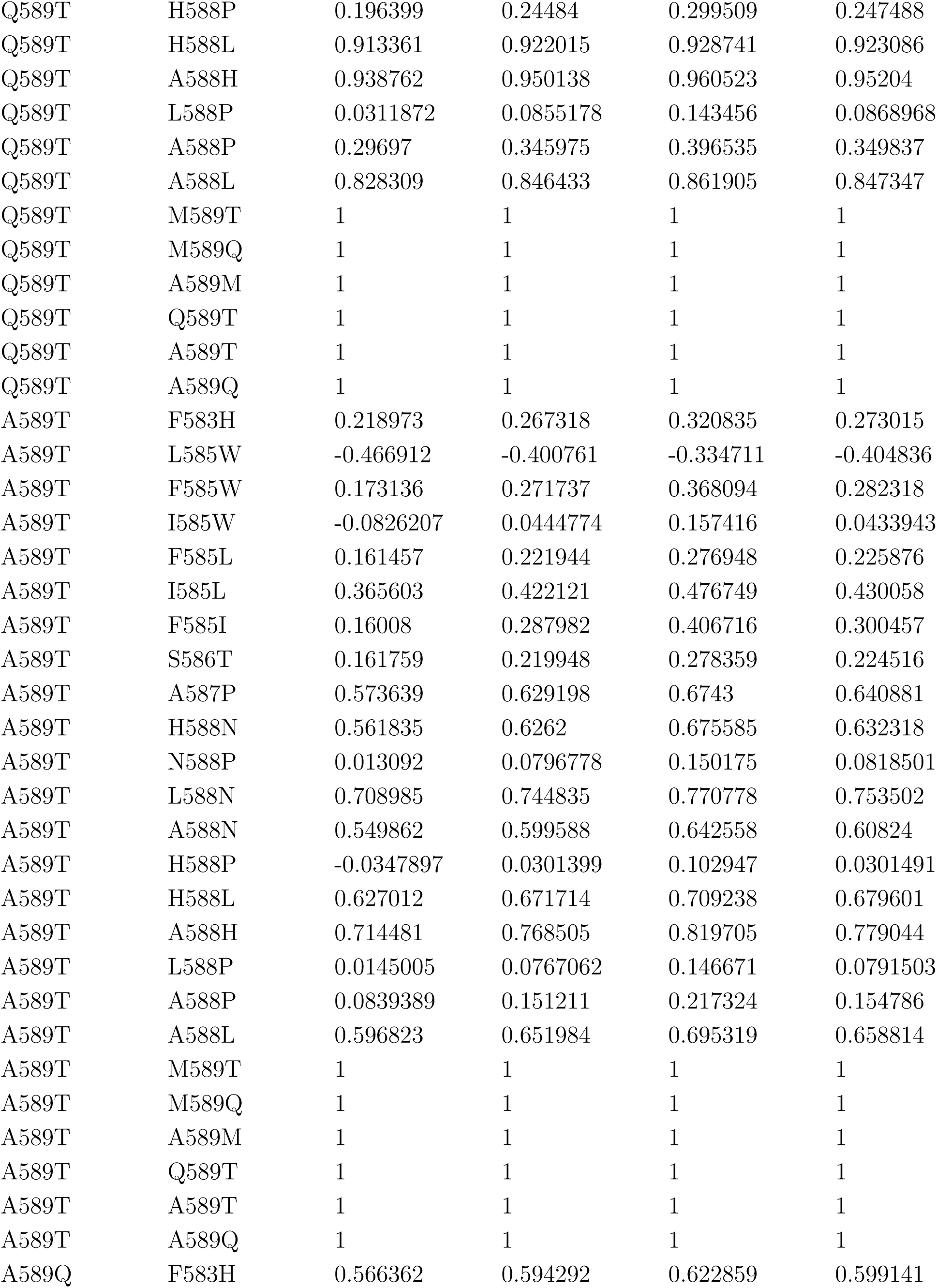
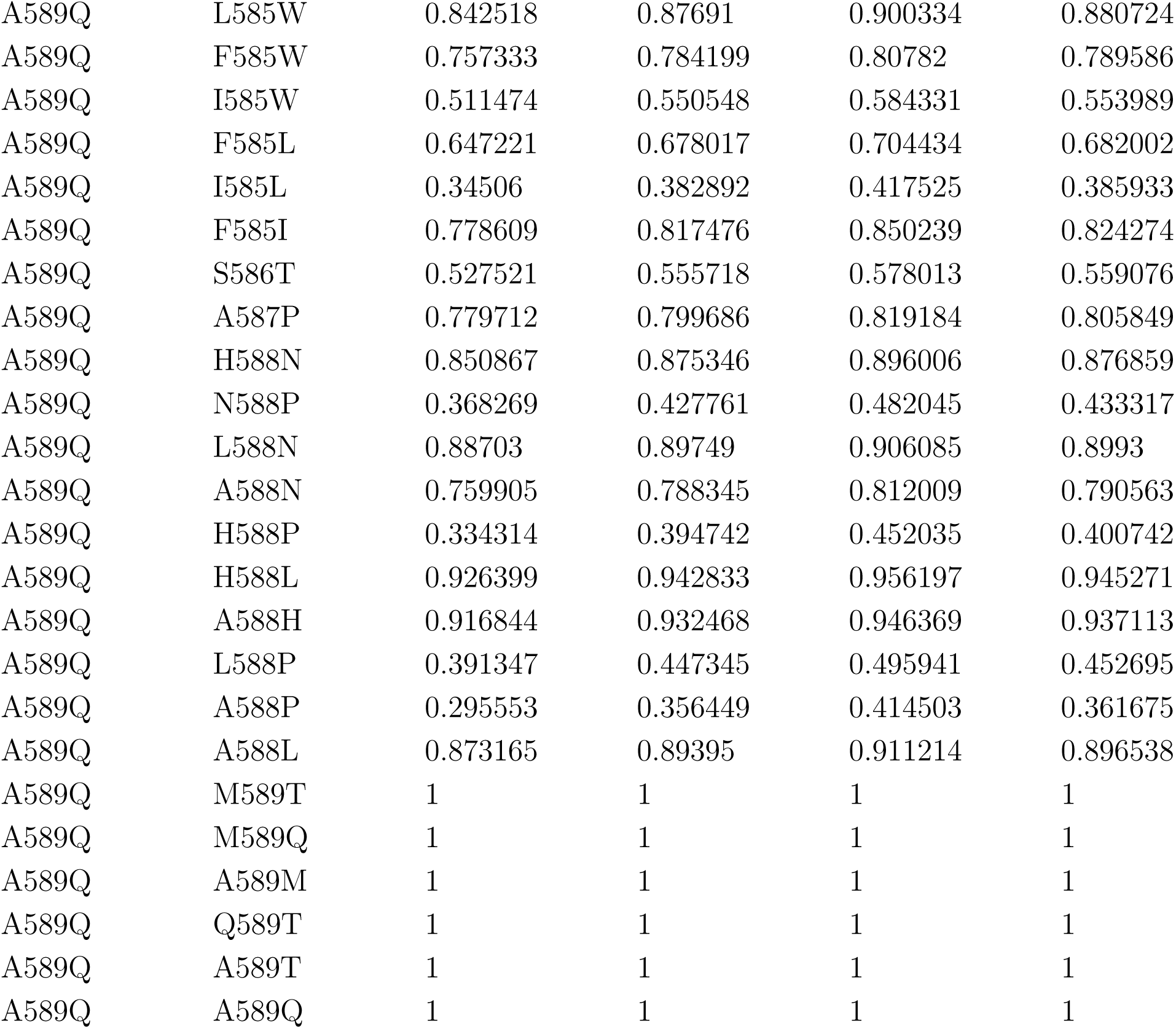
Epistasis measure 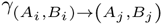 (eq. S1_8) between any two substitutions, averaged across the entire landscape calculated over 1,000 posterior samples. Shown are the 2.5%, 50%, 97.5% confidence intervals and the corresponding value calculated from the median growth rates. The majority of interactions are small to moderate.

| $A_i B_i$ | $A_j B_j$ | 2.5%C.I. | 50%C.I. | 97.5%C.I. | Main text |
| --- | --- | --- | --- | --- | --- |
| F583H | F583H | 1 | 1 | 1 | 1 |
| F583H | L585W | 0.781186 | 0.7957 | 0.808314 | 0.798262 |
| F583H | F585W | 0.656044 | 0.676274 | 0.695468 | 0.680326 |
| F583H | I585W | 0.429508 | 0.457521 | 0.482397 | 0.459904 |
| F583H | F585L | 0.777136 | 0.794864 | 0.812339 | 0.7982 |
| F583H | I585L | 0.569344 | 0.591398 | 0.612607 | 0.59463 |
| F583H | F585I | 0.773181 | 0.797178 | 0.81682 | 0.802641 |
| F583H | S586T | 0.693191 | 0.708917 | 0.724002 | 0.713048 |
| F583H | A587P | 0.754603 | 0.766899 | 0.779039 | 0.771487 |
| F583H | H588N | 0.842544 | 0.854133 | 0.864508 | 0.856289 |
| F583H | N588P | 0.190576 | 0.229818 | 0.269909 | 0.232421 |
| F583H | L588N | 0.925346 | 0.933872 | 0.941742 | 0.936301 |
| F583H | A588N | 0.763268 | 0.778255 | 0.790182 | 0.779749 |
| F583H | H588P | 0.233406 | 0.269719 | 0.310512 | 0.272072 |
| F583H | H588L | 0.897587 | 0.906249 | 0.913168 | 0.908253 |
| F583H | A588H | 0.935011 | 0.94343 | 0.95087 | 0.945953 |
| F583H | L588P | 0.164165 | 0.203443 | 0.244933 | 0.206263 |
| F583H | A588P | 0.302983 | 0.339205 | 0.378535 | 0.341537 |
| F583H | A588L | 0.829686 | 0.839552 | 0.848894 | 0.841511 |
| F583H | M589T | 0.36478 | 0.388584 | 0.410715 | 0.390504 |
| F583H | M589Q | 0.767691 | 0.787224 | 0.805853 | 0.791224 |
| F583H | A589M | 0.475742 | 0.501599 | 0.524351 | 0.505354 |
| F583H | Q589T | 0.561673 | 0.582565 | 0.599189 | 0.584278 |
| F583H | A589T | 0.719254 | 0.743653 | 0.766629 | 0.748196 |
| F583H | A589Q | 0.660341 | 0.68148 | 0.704744 | 0.685819 |
| L585W | F583H | 0.454182 | 0.487632 | 0.518399 | 0.491896 |
| L585W | L585W | 1 | 1 | 1 | 1 |
| L585W | F585W | 1 | 1 | 1 | 1 |
| L585W | I585W | 1 | 1 | 1 | 1 |
| L585W | F585L | 1 | 1 | 1 | 1 |
| L585W | I585L | 1 | 1 | 1 | 1 |
| L585W | F585I | 1 | 1 | 1 | 1 |
| L585W | S586T | 0.473765 | 0.514234 | 0.547457 | 0.517321 |
| L585W | A587P | 0.518897 | 0.554512 | 0.583423 | 0.559365 |
| L585W | H588N | 0.726887 | 0.743854 | 0.760751 | 0.746688 |
| L585W | N588P | -0.0963246 | -0.0408707 | 0.0188961 | -0.0403157 |
| L585W | L588N | 0.773517 | 0.797603 | 0.818312 | 0.799818 |
| L585W | A588N | 0.637654 | 0.658128 | 0.678935 | 0.660951 |
| L585W | H588P | -0.00537098 | 0.054329 | 0.114804 | 0.0547264 |
| L585W | H588L | 0.790838 | 0.807674 | 0.82184 | 0.80912 |
| L585W | A588H | 0.942444 | 0.954765 | 0.964412 | 0.95773 |
| L585W | L588P | -0.0629087 | -3.80E-05 | 0.0654933 | 0.000262741 |
| L585W | A588P | 0.0306047 | 0.0885322 | 0.153985 | 0.0907307 |
| L585W | A588L | 0.712324 | 0.737825 | 0.762529 | 0.740075 |
| L585W | M589T | -0.0422936 | -0.0160635 | 0.0106293 | -0.0159503 |
| L585W | M589Q | 0.182411 | 0.255928 | 0.320006 | 0.259131 |
| L585W | A589M | -0.310251 | -0.215613 | -0.123525 | -0.219263 |
| L585W | Q589T | -0.0942194 | -0.0693456 | -0.0443903 | -0.0696156 |
| L585W | A589T | -0.407908 | -0.361778 | -0.305441 | -0.363797 |
| L585W | A589Q | 0.275792 | 0.386399 | 0.479367 | 0.393986 |
| F585W | F583H | 0.575435 | 0.60054 | 0.624705 | 0.604693 |
| F585W | L585W | 1 | 1 | 1 | 1 |
| F585W | F585W | 1 | 1 | 1 | 1 |
| F585W | I585W | 1 | 1 | 1 | 1 |
| F585W | F585L | 1 | 1 | 1 | 1 |
| F585W | I585L | 1 | 1 | 1 | 1 |
| F585W | F585I | 1 | 1 | 1 | 1 |
| F585W | S586T | 0.563464 | 0.592818 | 0.617245 | 0.597256 |
| F585W | A587P | 0.63104 | 0.655021 | 0.675156 | 0.659723 |
| F585W | H588N | 0.849929 | 0.873172 | 0.890695 | 0.876316 |
| F585W | N588P | 0.0748116 | 0.124129 | 0.168615 | 0.126382 |
| F585W | L588N | 0.82937 | 0.853201 | 0.871674 | 0.855305 |
| F585W | A588N | 0.853412 | 0.86723 | 0.879036 | 0.87048 |
| F585W | H588P | 0.127938 | 0.175822 | 0.222216 | 0.178053 |
| F585W | H588L | 0.822823 | 0.842117 | 0.857598 | 0.844528 |
| F585W | A588H | 0.924808 | 0.941599 | 0.954542 | 0.944652 |
| F585W | L588P | 0.0147719 | 0.0596892 | 0.100846 | 0.0609615 |
| F585W | A588P | 0.125846 | 0.178084 | 0.222375 | 0.179708 |
| F585W | A588L | 0.770335 | 0.789736 | 0.806575 | 0.791355 |
| F585W | M589T | 0.420734 | 0.451691 | 0.48096 | 0.453511 |
| F585W | M589Q | 0.619419 | 0.664335 | 0.70375 | 0.667214 |
| F585W | A589M | 0.17335 | 0.259591 | 0.328658 | 0.262461 |
| F585W | Q589T | 0.618766 | 0.640466 | 0.659139 | 0.643393 |
| F585W | A589T | 0.72921 | 0.760218 | 0.788596 | 0.767587 |
| F585W | A589Q | 0.515032 | 0.570298 | 0.616933 | 0.576526 |
| I585W | F583H | 0.565929 | 0.585825 | 0.606133 | 0.588829 |
| I585W | L585W | 1 | 1 | 1 | 1 |
| I585W | F585W | 1 | 1 | 1 | 1 |
| I585W | I585W | 1 | 1 | 1 | 1 |
| I585W | F585L | 1 | 1 | 1 | 1 |
| I585W | I585L | 1 | 1 | 1 | 1 |
| I585W | F585I | 1 | 1 | 1 | 1 |
| I585W | S586T | 0.718259 | 0.74288 | 0.762948 | 0.746601 |
| I585W | A587P | 0.642102 | 0.661945 | 0.680028 | 0.66543 |
| I585W | H588N | 0.734221 | 0.749666 | 0.764296 | 0.751656 |
| I585W | N588P | -0.0953089 | -0.0524529 | -0.0117559 | -0.0529523 |
| I585W | L588N | 0.841038 | 0.854263 | 0.865488 | 0.856459 |
| I585W | A588N | 0.636411 | 0.655251 | 0.674846 | 0.657185 |
| I585W | H588P | 0.0813317 | 0.121835 | 0.163185 | 0.123579 |
| I585W | H588L | 0.907622 | 0.919595 | 0.928475 | 0.921124 |
| I585W | A588H | 0.943274 | 0.953209 | 0.960734 | 0.955383 |
| I585W | L588P | 0.007348 | 0.0529695 | 0.0958002 | 0.0534012 |
| I585W | A588P | 0.130452 | 0.173724 | 0.215956 | 0.17604 |
| I585W | A588L | 0.877774 | 0.88869 | 0.898551 | 0.890557 |
| I585W | M589T | 0.340237 | 0.366168 | 0.390183 | 0.367498 |
| I585W | M589Q | 0.67767 | 0.711942 | 0.742175 | 0.715303 |
| I585W | A589M | 0.196538 | 0.241079 | 0.287418 | 0.244517 |
| I585W | Q589T | 0.606682 | 0.621816 | 0.634968 | 0.623444 |
| I585W | A589T | 0.773898 | 0.799966 | 0.8209 | 0.804039 |
| I585W | A589Q | 0.509127 | 0.546985 | 0.583251 | 0.552331 |
| F585L | F583H | 0.498713 | 0.537351 | 0.578225 | 0.542494 |
| F585L | L585W | 1 | 1 | 1 | 1 |
| F585L | F585W | 1 | 1 | 1 | 1 |
| F585L | I585W | 1 | 1 | 1 | 1 |
| F585L | F585L | 1 | 1 | 1 | 1 |
| F585L | I585L | 1 | 1 | 1 | 1 |
| F585L | F585I | 1 | 1 | 1 | 1 |
| F585L | S586T | 0.508648 | 0.544855 | 0.577364 | 0.550048 |
| F585L | A587P | 0.748831 | 0.771728 | 0.792241 | 0.78029 |
| F585L | H588N | 0.783446 | 0.81465 | 0.838701 | 0.818782 |
| F585L | N588P | 0.174674 | 0.231065 | 0.283916 | 0.234195 |
| F585L | L588N | 0.842643 | 0.859868 | 0.873231 | 0.864198 |
| F585L | A588N | 0.758322 | 0.781194 | 0.802305 | 0.786606 |
| F585L | H588P | 0.234263 | 0.287353 | 0.344957 | 0.290631 |
| F585L | H588L | 0.867885 | 0.893601 | 0.912608 | 0.898159 |
| F585L | A588H | 0.902161 | 0.926496 | 0.944877 | 0.932376 |
| F585L | L588P | 0.119219 | 0.174994 | 0.225747 | 0.177166 |
| F585L | A588P | 0.239477 | 0.293336 | 0.340646 | 0.297365 |
| F585L | A588L | 0.867397 | 0.883835 | 0.897475 | 0.888359 |
| F585L | M589T | 0.222748 | 0.324079 | 0.417104 | 0.331449 |
| F585L | M589Q | 0.731643 | 0.775209 | 0.807377 | 0.77954 |
| F585L | A589M | 0.508923 | 0.538641 | 0.566798 | 0.544187 |
| F585L | Q589T | 0.014926 | 0.0878362 | 0.16612 | 0.0910583 |
| F585L | A589T | 0.136648 | 0.183999 | 0.228006 | 0.186528 |
| F585L | A589Q | 0.617723 | 0.654863 | 0.683122 | 0.658132 |
| I585L | F583H | 0.440601 | 0.466005 | 0.493341 | 0.469596 |
| I585L | L585W | 1 | 1 | 1 | 1 |
| I585L | F585W | 1 | 1 | 1 | 1 |
| I585L | I585W | 1 | 1 | 1 | 1 |
| I585L | F585L | 1 | 1 | 1 | 1 |
| I585L | I585L | 1 | 1 | 1 | 1 |
| I585L | F585I | 1 | 1 | 1 | 1 |
| I585L | S586T | 0.731721 | 0.751609 | 0.772371 | 0.757651 |
| I585L | A587P | 0.722903 | 0.740117 | 0.755216 | 0.745156 |
| I585L | H588N | 0.703994 | 0.725282 | 0.746375 | 0.728149 |
| I585L | N588P | -0.0147815 | 0.0390647 | 0.0912781 | 0.0384516 |
| I585L | L588N | 0.780189 | 0.798246 | 0.813711 | 0.801102 |
| I585L | A588N | 0.601557 | 0.627081 | 0.650434 | 0.629603 |
| I585L | H588P | 0.131625 | 0.184265 | 0.236204 | 0.185243 |
| I585L | H588L | 0.897078 | 0.912996 | 0.923876 | 0.91505 |
| I585L | A588H | 0.916351 | 0.932322 | 0.946543 | 0.936478 |
| I585L | L588P | 0.0589946 | 0.118835 | 0.171694 | 0.118318 |
| I585L | A588P | 0.192648 | 0.24165 | 0.290474 | 0.244484 |
| I585L | A588L | 0.832471 | 0.851941 | 0.868587 | 0.854602 |
| I585L | M589T | 0.326097 | 0.388791 | 0.443753 | 0.393248 |
| I585L | M589Q | 0.713169 | 0.744019 | 0.772596 | 0.748478 |
| I585L | A589M | 0.414194 | 0.439381 | 0.461654 | 0.441654 |
| I585L | Q589T | 0.378748 | 0.425656 | 0.466665 | 0.427833 |
| I585L | A589T | 0.628706 | 0.659398 | 0.688126 | 0.665311 |
| I585L | A589Q | 0.580186 | 0.605879 | 0.630692 | 0.609325 |
| F585I | F583H | 0.184445 | 0.249108 | 0.306907 | 0.255675 |
| F585I | L585W | 1 | 1 | 1 | 1 |
| F585I | F585W | 1 | 1 | 1 | 1 |
| F585I | I585W | 1 | 1 | 1 | 1 |
| F585I | F585L | 1 | 1 | 1 | 1 |
| F585I | I585L | 1 | 1 | 1 | 1 |
| F585I | F585I | 1 | 1 | 1 | 1 |
| F585I | S586T | 0.375036 | 0.441937 | 0.506261 | 0.45574 |
| F585I | A587P | 0.237367 | 0.294618 | 0.348139 | 0.302799 |
| F585I | H588N | 0.0905469 | 0.163255 | 0.226801 | 0.162977 |
| F585I | N588P | -0.207737 | -0.119257 | -0.0320937 | -0.124733 |
| F585I | L588N | 0.296632 | 0.3553 | 0.409815 | 0.361256 |
| F585I | A588N | 0.0843689 | 0.148095 | 0.212808 | 0.150577 |
| F585I | H588P | -0.0721975 | 0.0221204 | 0.123655 | 0.0217359 |
| F585I | H588L | 0.682545 | 0.739801 | 0.786965 | 0.748623 |
| F585I | A588H | 0.746857 | 0.80704 | 0.857383 | 0.820367 |
| F585I | L588P | -0.244073 | -0.134591 | -0.0286072 | -0.141615 |
| F585I | A588P | -0.0773802 | 0.0199233 | 0.130122 | 0.0208728 |
| F585I | A588L | 0.679159 | 0.739456 | 0.788322 | 0.746859 |
| F585I | M589T | -0.0223809 | 0.0369155 | 0.0939654 | 0.0384758 |
| F585I | M589Q | 0.181643 | 0.304356 | 0.438138 | 0.316988 |
| F585I | A589M | 0.115825 | 0.207888 | 0.302134 | 0.219138 |
| F585I | Q589T | 0.258418 | 0.331378 | 0.392724 | 0.337253 |
| F585I | A589T | 0.408696 | 0.498139 | 0.575123 | 0.512365 |
| F585I | A589Q | 0.192987 | 0.305332 | 0.413546 | 0.314907 |
| S586T | F583H | 0.558533 | 0.578932 | 0.600615 | 0.583796 |
| S586T | L585W | 0.668146 | 0.692647 | 0.715473 | 0.695892 |
| S586T | F585W | 0.60444 | 0.629865 | 0.653122 | 0.634002 |
| S586T | I585W | 0.405514 | 0.450451 | 0.493272 | 0.454801 |
| S586T | F585L | 0.715555 | 0.737153 | 0.757711 | 0.74183 |
| S586T | I585L | 0.626532 | 0.653608 | 0.682645 | 0.661327 |
| S586T | F585I | 0.76627 | 0.796508 | 0.823164 | 0.80464 |
| S586T | S586T | 1 | 1 | 1 | 1 |
| S586T | A587P | 0.756743 | 0.777474 | 0.79646 | 0.784579 |
| S586T | H588N | 0.757426 | 0.77784 | 0.794093 | 0.781394 |
| S586T | N588P | 0.117798 | 0.163729 | 0.20692 | 0.166125 |
| S586T | L588N | 0.803608 | 0.814496 | 0.826797 | 0.81762 |
| S586T | A588N | 0.669584 | 0.692896 | 0.714201 | 0.695518 |
| S586T | H588P | 0.183005 | 0.226778 | 0.271477 | 0.229452 |
| S586T | H588L | 0.905321 | 0.919734 | 0.931172 | 0.922732 |
| S586T | A588H | 0.895396 | 0.912135 | 0.925907 | 0.916249 |
| S586T | L588P | 0.156165 | 0.200199 | 0.244447 | 0.202201 |
| S586T | A588P | 0.210221 | 0.252071 | 0.299017 | 0.25654 |
| S586T | A588L | 0.848712 | 0.864269 | 0.877965 | 0.866903 |
| S586T | M589T | 0.320181 | 0.34948 | 0.376496 | 0.350684 |
| S586T | M589Q | 0.444158 | 0.500164 | 0.546745 | 0.503497 |
| S586T | A589M | 0.501336 | 0.528305 | 0.555114 | 0.53348 |
| S586T | Q589T | 0.283318 | 0.304245 | 0.325068 | 0.306711 |
| S586T | A589T | 0.657295 | 0.686278 | 0.711797 | 0.690561 |
| S586T | A589Q | 0.330932 | 0.363657 | 0.390418 | 0.366934 |
| A587P | F583H | 0.444399 | 0.471354 | 0.495931 | 0.477261 |
| A587P | L585W | 0.458245 | 0.500674 | 0.532394 | 0.504322 |
| A587P | F585W | 0.40003 | 0.441218 | 0.477764 | 0.445334 |
| A587P | I585W | 0.132874 | 0.177005 | 0.216795 | 0.178225 |
| A587P | F585L | 0.674785 | 0.70362 | 0.727819 | 0.714168 |
| A587P | I585L | 0.472926 | 0.500006 | 0.52672 | 0.507174 |
| A587P | F585I | 0.591916 | 0.626418 | 0.658595 | 0.635448 |
| A587P | S586T | 0.619421 | 0.650491 | 0.681307 | 0.660246 |
| A587P | A587P | 1 | 1 | 1 | 1 |
| A587P | H588N | 0.638158 | 0.671336 | 0.701392 | 0.673741 |
| A587P | N588P | 0.153513 | 0.207187 | 0.252698 | 0.209571 |
| A587P | L588N | 0.766936 | 0.785239 | 0.802774 | 0.78839 |
| A587P | A588N | 0.62863 | 0.661831 | 0.688638 | 0.663819 |
| A587P | H588P | 0.106974 | 0.152807 | 0.191954 | 0.154351 |
| A587P | H588L | 0.711105 | 0.747364 | 0.778275 | 0.752304 |
| A587P | A588H | 0.830963 | 0.862226 | 0.887217 | 0.870824 |
| A587P | L588P | -0.0440905 | 0.00351999 | 0.0467686 | 0.00313926 |
| A587P | A588P | 0.0369484 | 0.0817756 | 0.127242 | 0.0829486 |
| A587P | A588L | 0.679179 | 0.709999 | 0.735459 | 0.715062 |
| A587P | M589T | 0.214566 | 0.254008 | 0.290491 | 0.25677 |
| A587P | M589Q | 0.558199 | 0.594929 | 0.625845 | 0.600397 |
| A587P | A589M | 0.269509 | 0.317502 | 0.357892 | 0.322202 |
| A587P | Q589T | 0.525878 | 0.554338 | 0.580558 | 0.560294 |
| A587P | A589T | 0.652057 | 0.692982 | 0.728145 | 0.703486 |
| A587P | A589Q | 0.584218 | 0.623677 | 0.65867 | 0.632985 |
| H588N | F583H | 0.364167 | 0.40803 | 0.447776 | 0.411761 |
| H588N | L585W | 0.305853 | 0.347263 | 0.385203 | 0.350732 |
| H588N | F585W | 0.388132 | 0.45727 | 0.513089 | 0.464171 |
| H588N | I585W | 0.119186 | 0.155796 | 0.188542 | 0.157692 |
| H588N | F585L | 0.561127 | 0.615694 | 0.661239 | 0.622816 |
| H588N | I585L | 0.41906 | 0.450719 | 0.480376 | 0.454573 |
| H588N | F585I | 0.421243 | 0.477581 | 0.525674 | 0.481796 |
| H588N | S586T | 0.459634 | 0.498636 | 0.533117 | 0.504342 |
| H588N | A587P | 0.326995 | 0.386713 | 0.441572 | 0.388548 |
| H588N | H588N | 1 | 1 | 1 | 1 |
| H588N | N588P | 1 | 1 | 1 | 1 |
| H588N | L588N | 1 | 1 | 1 | 1 |
| H588N | A588N | 1 | 1 | 1 | 1 |
| H588N | H588P | 1 | 1 | 1 | 1 |
| H588N | H588L | 1 | 1 | 1 | 1 |
| H588N | A588H | 1 | 1 | 1 | 1 |
| H588N | L588P | 1 | 1 | 1 | 1 |
| H588N | A588P | 1 | 1 | 1 | 1 |
| H588N | A588L | 1 | 1 | 1 | 1 |
| H588N | M589T | 0.1649 | 0.214481 | 0.262116 | 0.214107 |
| H588N | M589Q | 0.646526 | 0.68125 | 0.711678 | 0.685033 |
| H588N | A589M | 0.260622 | 0.36808 | 0.459375 | 0.370694 |
| H588N | Q589T | 0.355734 | 0.402642 | 0.446682 | 0.404114 |
| H588N | A589T | 0.541341 | 0.613113 | 0.66742 | 0.619917 |
| H588N | A589Q | 0.372683 | 0.471628 | 0.558053 | 0.475109 |
| N588P | F583H | 0.796886 | 0.807261 | 0.817995 | 0.809733 |
| N588P | L585W | 0.864462 | 0.874946 | 0.88532 | 0.876739 |
| N588P | F585W | 0.780449 | 0.798435 | 0.811425 | 0.800614 |
| N588P | I585W | 0.658127 | 0.676268 | 0.692689 | 0.678165 |
| N588P | F585L | 0.87595 | 0.885417 | 0.894828 | 0.88751 |
| N588P | I585L | 0.800502 | 0.812417 | 0.823228 | 0.814426 |
| N588P | F585I | 0.921288 | 0.931104 | 0.940546 | 0.934299 |
| N588P | S586T | 0.8529 | 0.862946 | 0.870969 | 0.864982 |
| N588P | A587P | 0.852285 | 0.862262 | 0.870824 | 0.86468 |
| N588P | H588N | 1 | 1 | 1 | 1 |
| N588P | N588P | 1 | 1 | 1 | 1 |
| N588P | L588N | 1 | 1 | 1 | 1 |
| N588P | A588N | 1 | 1 | 1 | 1 |
| N588P | H588P | 1 | 1 | 1 | 1 |
| N588P | H588L | 1 | 1 | 1 | 1 |
| N588P | A588H | 1 | 1 | 1 | 1 |
| N588P | L588P | 1 | 1 | 1 | 1 |
| N588P | A588P | 1 | 1 | 1 | 1 |
| N588P | A588L | 1 | 1 | 1 | 1 |
| N588P | M589T | 0.558056 | 0.58033 | 0.603979 | 0.582521 |
| N588P | M589Q | 0.88279 | 0.893628 | 0.903445 | 0.895108 |
| N588P | A589M | 0.733958 | 0.755138 | 0.775567 | 0.757921 |
| N588P | Q589T | 0.746034 | 0.762406 | 0.78065 | 0.76479 |
| N588P | A589T | 0.83446 | 0.852405 | 0.8691 | 0.855846 |
| N588P | A589Q | 0.857413 | 0.871084 | 0.882708 | 0.873541 |
| L588N | F583H | 0.572777 | 0.623287 | 0.667224 | 0.632426 |
| L588N | L585W | 0.496956 | 0.543582 | 0.586391 | 0.546829 |
| L588N | F585W | 0.55716 | 0.612614 | 0.658266 | 0.615578 |
| L588N | I585W | 0.169691 | 0.231451 | 0.289624 | 0.235697 |
| L588N | F585L | 0.585495 | 0.624308 | 0.657367 | 0.634131 |
| L588N | I585L | -0.128987 | -0.0709644 | -0.00350338 | -0.0698335 |
| L588N | F585I | 0.0869328 | 0.149603 | 0.210072 | 0.15148 |
| L588N | S586T | 0.163505 | 0.207983 | 0.252605 | 0.213155 |
| L588N | A587P | 0.213638 | 0.269562 | 0.323857 | 0.273392 |
| L588N | H588N | 1 | 1 | 1 | 1 |
| L588N | N588P | 1 | 1 | 1 | 1 |
| L588N | L588N | 1 | 1 | 1 | 1 |
| L588N | A588N | 1 | 1 | 1 | 1 |
| L588N | H588P | 1 | 1 | 1 | 1 |
| L588N | H588L | 1 | 1 | 1 | 1 |
| L588N | A588H | 1 | 1 | 1 | 1 |
| L588N | L588P | 1 | 1 | 1 | 1 |
| L588N | A588P | 1 | 1 | 1 | 1 |
| L588N | A588L | 1 | 1 | 1 | 1 |
| L588N | M589T | 0.473429 | 0.531185 | 0.578612 | 0.536087 |
| L588N | M589Q | 0.634864 | 0.672167 | 0.706915 | 0.677271 |
| L588N | A589M | 0.420305 | 0.47321 | 0.518072 | 0.479182 |
| L588N | Q589T | 0.317629 | 0.374117 | 0.419175 | 0.378957 |
| L588N | A589T | 0.640882 | 0.684853 | 0.717536 | 0.694917 |
| L588N | A589Q | 0.266508 | 0.310993 | 0.350187 | 0.314844 |
| A588N | F583H | 0.276645 | 0.314086 | 0.349309 | 0.315059 |
| A588N | L585W | 0.420779 | 0.452762 | 0.486716 | 0.45531 |
| A588N | F585W | 0.529207 | 0.577347 | 0.613812 | 0.582731 |
| A588N | I585W | 0.0747905 | 0.104877 | 0.13589 | 0.105309 |
| A588N | F585L | 0.652158 | 0.687974 | 0.720333 | 0.694435 |
| A588N | I585L | 0.41337 | 0.443434 | 0.470715 | 0.445503 |
| A588N | F585I | 0.569462 | 0.60663 | 0.637953 | 0.611501 |
| A588N | S586T | 0.445508 | 0.478259 | 0.509802 | 0.481663 |
| A588N | A587P | 0.50433 | 0.545568 | 0.584122 | 0.548711 |
| A588N | H588N | 1 | 1 | 1 | 1 |
| A588N | N588P | 1 | 1 | 1 | 1 |
| A588N | L588N | 1 | 1 | 1 | 1 |
| A588N | A588N | 1 | 1 | 1 | 1 |
| A588N | H588P | 1 | 1 | 1 | 1 |
| A588N | H588L | 1 | 1 | 1 | 1 |
| A588N | A588H | 1 | 1 | 1 | 1 |
| A588N | L588P | 1 | 1 | 1 | 1 |
| A588N | A588P | 1 | 1 | 1 | 1 |
| A588N | A588L | 1 | 1 | 1 | 1 |
| A588N | M589T | 0.0758398 | 0.140718 | 0.197795 | 0.138998 |
| A588N | M589Q | 0.68905 | 0.718399 | 0.744794 | 0.721058 |
| A588N | A589M | 0.242384 | 0.309005 | 0.375651 | 0.310863 |
| A588N | Q589T | 0.242963 | 0.302766 | 0.35675 | 0.302634 |
| A588N | A589T | 0.690495 | 0.729725 | 0.759972 | 0.737069 |
| A588N | A589Q | 0.450311 | 0.51089 | 0.562515 | 0.514914 |
| H588P | F583H | 0.722663 | 0.736607 | 0.751248 | 0.739238 |
| H588P | L585W | 0.833589 | 0.846533 | 0.857924 | 0.848363 |
| H588P | F585W | 0.730656 | 0.74906 | 0.764905 | 0.751802 |
| H588P | I585W | 0.503165 | 0.527783 | 0.551007 | 0.52969 |
| H588P | F585L | 0.846354 | 0.85907 | 0.871503 | 0.861465 |
| H588P | I585L | 0.696712 | 0.716426 | 0.735834 | 0.719191 |
| H588P | F585I | 0.873775 | 0.889833 | 0.904735 | 0.89347 |
| H588P | S586T | 0.79044 | 0.80333 | 0.815052 | 0.806012 |
| H588P | A587P | 0.852984 | 0.861531 | 0.868901 | 0.864671 |
| H588P | H588N | 1 | 1 | 1 | 1 |
| H588P | N588P | 1 | 1 | 1 | 1 |
| H588P | L588N | 1 | 1 | 1 | 1 |
| H588P | A588N | 1 | 1 | 1 | 1 |
| H588P | H588P | 1 | 1 | 1 | 1 |
| H588P | H588L | 1 | 1 | 1 | 1 |
| H588P | A588H | 1 | 1 | 1 | 1 |
| H588P | L588P | 1 | 1 | 1 | 1 |
| H588P | A588P | 1 | 1 | 1 | 1 |
| H588P | A588L | 1 | 1 | 1 | 1 |
| H588P | M589T | 0.428294 | 0.457786 | 0.488056 | 0.460445 |
| H588P | M589Q | 0.859507 | 0.872401 | 0.88332 | 0.874244 |
| H588P | A589M | 0.67024 | 0.696139 | 0.718054 | 0.699549 |
| H588P | Q589T | 0.675904 | 0.697057 | 0.718344 | 0.699143 |
| H588P | A589T | 0.735399 | 0.760289 | 0.783452 | 0.763774 |
| H588P | A589Q | 0.824131 | 0.840084 | 0.855654 | 0.843174 |
| H588L | F583H | 0.537322 | 0.570348 | 0.59877 | 0.575948 |
| H588L | L585W | 0.483839 | 0.522418 | 0.557248 | 0.52512 |
| H588L | F585W | 0.505674 | 0.548758 | 0.58806 | 0.552972 |
| H588L | I585W | 0.322656 | 0.387631 | 0.440227 | 0.391864 |
| H588L | F585L | 0.798026 | 0.837371 | 0.865515 | 0.843904 |
| H588L | I585L | 0.573102 | 0.631645 | 0.676372 | 0.637841 |
| H588L | F585I | 0.637739 | 0.701911 | 0.754867 | 0.710492 |
| H588L | S586T | 0.659783 | 0.709983 | 0.751646 | 0.718148 |
| H588L | A587P | 0.573428 | 0.628092 | 0.673632 | 0.633629 |
| H588L | H588N | 1 | 1 | 1 | 1 |
| H588L | N588P | 1 | 1 | 1 | 1 |
| H588L | L588N | 1 | 1 | 1 | 1 |
| H588L | A588N | 1 | 1 | 1 | 1 |
| H588L | H588P | 1 | 1 | 1 | 1 |
| H588L | H588L | 1 | 1 | 1 | 1 |
| H588L | A588H | 1 | 1 | 1 | 1 |
| H588L | L588P | 1 | 1 | 1 | 1 |
| H588L | A588P | 1 | 1 | 1 | 1 |
| H588L | A588L | 1 | 1 | 1 | 1 |
| H588L | M589T | 0.355372 | 0.402987 | 0.450246 | 0.406109 |
| H588L | M589Q | 0.61995 | 0.653075 | 0.682368 | 0.656049 |
| H588L | A589M | 0.30928 | 0.422855 | 0.529679 | 0.42886 |
| H588L | Q589T | 0.57079 | 0.608733 | 0.639655 | 0.612123 |
| H588L | A589T | 0.490726 | 0.551847 | 0.605044 | 0.561098 |
| H588L | A589Q | 0.599976 | 0.690501 | 0.763127 | 0.699435 |
| A588H | F583H | -0.121131 | -0.0311957 | 0.0587075 | -0.0342472 |
| A588H | L585W | 0.461349 | 0.554032 | 0.637626 | 0.571732 |
| A588H | F585W | 0.125621 | 0.262657 | 0.385727 | 0.271563 |
| A588H | I585W | -0.228101 | -0.123826 | -0.0220694 | -0.132762 |
| A588H | F585L | 0.421178 | 0.540025 | 0.645104 | 0.563555 |
| A588H | I585L | -0.0200111 | 0.113126 | 0.24589 | 0.121353 |
| A588H | F585I | 0.157248 | 0.327582 | 0.490841 | 0.349158 |
| A588H | S586T | -0.098815 | 0.0239002 | 0.140459 | 0.0252088 |
| A588H | A587P | 0.256828 | 0.375819 | 0.481963 | 0.39213 |
| A588H | H588N | 1 | 1 | 1 | 1 |
| A588H | N588P | 1 | 1 | 1 | 1 |
| A588H | L588N | 1 | 1 | 1 | 1 |
| A588H | A588N | 1 | 1 | 1 | 1 |
| A588H | H588P | 1 | 1 | 1 | 1 |
| A588H | H588L | 1 | 1 | 1 | 1 |
| A588H | A588H | 1 | 1 | 1 | 1 |
| A588H | L588P | 1 | 1 | 1 | 1 |
| A588H | A588P | 1 | 1 | 1 | 1 |
| A588H | A588L | 1 | 1 | 1 | 1 |
| A588H | M589T | -0.287865 | -0.152514 | -0.00945207 | -0.164418 |
| A588H | M589Q | 0.524019 | 0.600639 | 0.667946 | 0.61576 |
| A588H | A589M | -0.0516285 | 0.0519124 | 0.14294 | 0.0522902 |
| A588H | Q589T | -0.455615 | -0.328814 | -0.172035 | -0.346725 |
| A588H | A589T | -0.162622 | 0.00386973 | 0.176045 | 0.00539306 |
| A588H | A589Q | 0.283386 | 0.397965 | 0.50059 | 0.415179 |
| L588P | F583H | 0.815504 | 0.824543 | 0.833441 | 0.826696 |
| L588P | L585W | 0.879053 | 0.887793 | 0.896158 | 0.889284 |
| L588P | F585W | 0.809015 | 0.822135 | 0.832573 | 0.824236 |
| L588P | I585W | 0.629276 | 0.648119 | 0.666832 | 0.650521 |
| L588P | F585L | 0.885131 | 0.894142 | 0.902543 | 0.89599 |
| L588P | I585L | 0.762746 | 0.77789 | 0.791381 | 0.779594 |
| L588P | F585I | 0.897556 | 0.9096 | 0.920582 | 0.912231 |
| L588P | S586T | 0.832312 | 0.841767 | 0.851157 | 0.843883 |
| L588P | A587P | 0.869341 | 0.876453 | 0.883001 | 0.878686 |
| L588P | H588N | 1 | 1 | 1 | 1 |
| L588P | N588P | 1 | 1 | 1 | 1 |
| L588P | L588N | 1 | 1 | 1 | 1 |
| L588P | A588N | 1 | 1 | 1 | 1 |
| L588P | H588P | 1 | 1 | 1 | 1 |
| L588P | H588L | 1 | 1 | 1 | 1 |
| L588P | A588H | 1 | 1 | 1 | 1 |
| L588P | L588P | 1 | 1 | 1 | 1 |
| L588P | A588P | 1 | 1 | 1 | 1 |
| L588P | A588L | 1 | 1 | 1 | 1 |
| L588P | M589T | 0.615462 | 0.635788 | 0.655433 | 0.637349 |
| L588P | M589Q | 0.891889 | 0.901547 | 0.910663 | 0.903 |
| L588P | A589M | 0.753209 | 0.774391 | 0.792867 | 0.776988 |
| L588P | Q589T | 0.778945 | 0.79193 | 0.805796 | 0.793852 |
| L588P | A589T | 0.844111 | 0.859023 | 0.872826 | 0.862033 |
| L588P | A589Q | 0.865193 | 0.877239 | 0.888757 | 0.879972 |
| A588P | F583H | 0.708604 | 0.723912 | 0.740762 | 0.726943 |
| A588P | L585W | 0.821464 | 0.83457 | 0.847312 | 0.836936 |
| A588P | F585W | 0.71429 | 0.735461 | 0.753399 | 0.738473 |
| A588P | I585W | 0.47634 | 0.504271 | 0.53045 | 0.505953 |
| A588P | F585L | 0.833113 | 0.84668 | 0.857802 | 0.849264 |
| A588P | I585L | 0.667854 | 0.689099 | 0.709141 | 0.692123 |
| A588P | F585I | 0.86272 | 0.879768 | 0.896084 | 0.884655 |
| A588P | S586T | 0.771306 | 0.784702 | 0.798082 | 0.788293 |
| A588P | A587P | 0.825476 | 0.835171 | 0.844167 | 0.838456 |
| A588P | H588N | 1 | 1 | 1 | 1 |
| A588P | N588P | 1 | 1 | 1 | 1 |
| A588P | L588N | 1 | 1 | 1 | 1 |
| A588P | A588N | 1 | 1 | 1 | 1 |
| A588P | H588P | 1 | 1 | 1 | 1 |
| A588P | H588L | 1 | 1 | 1 | 1 |
| A588P | A588H | 1 | 1 | 1 | 1 |
| A588P | L588P | 1 | 1 | 1 | 1 |
| A588P | A588P | 1 | 1 | 1 | 1 |
| A588P | A588L | 1 | 1 | 1 | 1 |
| A588P | M589T | 0.519883 | 0.547342 | 0.574208 | 0.550427 |
| A588P | M589Q | 0.839046 | 0.854109 | 0.867391 | 0.856051 |
| A588P | A589M | 0.597928 | 0.627067 | 0.653776 | 0.630781 |
| A588P | Q589T | 0.748195 | 0.76731 | 0.784661 | 0.770048 |
| A588P | A589T | 0.765165 | 0.789067 | 0.810309 | 0.793702 |
| A588P | A589Q | 0.778221 | 0.796749 | 0.816004 | 0.80059 |
| A588L | F583H | 0.424192 | 0.454056 | 0.481585 | 0.457444 |
| A588L | L585W | 0.523416 | 0.56267 | 0.598961 | 0.564757 |
| A588L | F585W | 0.463087 | 0.504819 | 0.539237 | 0.507222 |
| A588L | I585W | 0.335864 | 0.378697 | 0.42374 | 0.382689 |
| A588L | F585L | 0.860671 | 0.878812 | 0.894559 | 0.883591 |
| A588L | I585L | 0.546684 | 0.597053 | 0.641602 | 0.602019 |
| A588L | F585I | 0.737578 | 0.789117 | 0.827323 | 0.795144 |
| A588L | S586T | 0.608846 | 0.646751 | 0.682133 | 0.65238 |
| A588L | A587P | 0.664308 | 0.697305 | 0.726469 | 0.70294 |
| A588L | H588N | 1 | 1 | 1 | 1 |
| A588L | N588P | 1 | 1 | 1 | 1 |
| A588L | L588N | 1 | 1 | 1 | 1 |
| A588L | A588N | 1 | 1 | 1 | 1 |
| A588L | H588P | 1 | 1 | 1 | 1 |
| A588L | H588L | 1 | 1 | 1 | 1 |
| A588L | A588H | 1 | 1 | 1 | 1 |
| A588L | L588P | 1 | 1 | 1 | 1 |
| A588L | A588P | 1 | 1 | 1 | 1 |
| A588L | A588L | 1 | 1 | 1 | 1 |
| A588L | M589T | 0.228406 | 0.289766 | 0.344052 | 0.28894 |
| A588L | M589Q | 0.726248 | 0.753999 | 0.778689 | 0.756972 |
| A588L | A589M | 0.407792 | 0.481079 | 0.544559 | 0.485841 |
| A588L | Q589T | 0.424406 | 0.477608 | 0.525877 | 0.479107 |
| A588L | A589T | 0.646349 | 0.697867 | 0.73463 | 0.704303 |
| A588L | A589Q | 0.648952 | 0.706927 | 0.755811 | 0.712652 |
| M589T | F583H | 0.650735 | 0.664927 | 0.677971 | 0.666804 |
| M589T | L585W | 0.54703 | 0.56318 | 0.577438 | 0.564 |
| M589T | F585W | 0.480907 | 0.51288 | 0.539057 | 0.515301 |
| M589T | I585W | 0.287422 | 0.316458 | 0.341994 | 0.317671 |
| M589T | F585L | 0.911228 | 0.922846 | 0.933026 | 0.92536 |
| M589T | I585L | 0.853407 | 0.868474 | 0.880963 | 0.870945 |
| M589T | F585I | 0.826379 | 0.840009 | 0.851789 | 0.843199 |
| M589T | S586T | 0.671977 | 0.688704 | 0.70288 | 0.690386 |
| M589T | A587P | 0.796758 | 0.808119 | 0.819254 | 0.810731 |
| M589T | H588N | 0.863383 | 0.874391 | 0.883686 | 0.875236 |
| M589T | N588P | -0.0778001 | -0.0268903 | 0.0207394 | -0.026676 |
| M589T | L588N | 0.949011 | 0.95541 | 0.960753 | 0.9563 |
| M589T | A588N | 0.826234 | 0.840939 | 0.853505 | 0.841498 |
| M589T | H588P | 0.0750677 | 0.119964 | 0.165821 | 0.122608 |
| M589T | H588L | 0.905195 | 0.914454 | 0.921855 | 0.91537 |
| M589T | A588H | 0.970463 | 0.977123 | 0.982399 | 0.978197 |
| M589T | L588P | -0.0777429 | -0.0284708 | 0.0193453 | -0.0279023 |
| M589T | A588P | 0.180031 | 0.223405 | 0.266025 | 0.225898 |
| M589T | A588L | 0.867986 | 0.881186 | 0.892501 | 0.881907 |
| M589T | M589T | 1 | 1 | 1 | 1 |
| M589T | M589Q | 1 | 1 | 1 | 1 |
| M589T | A589M | 1 | 1 | 1 | 1 |
| M589T | Q589T | 1 | 1 | 1 | 1 |
| M589T | A589T | 1 | 1 | 1 | 1 |
| M589T | A589Q | 1 | 1 | 1 | 1 |
| M589Q | F583H | 0.276666 | 0.331453 | 0.388945 | 0.338601 |
| M589Q | L585W | 0.366534 | 0.423993 | 0.47329 | 0.427614 |
| M589Q | F585W | 0.230867 | 0.3144 | 0.389001 | 0.318897 |
| M589Q | I585W | 0.0347747 | 0.11596 | 0.200697 | 0.116472 |
| M589Q | F585L | 0.423183 | 0.502226 | 0.568661 | 0.509794 |
| M589Q | I585L | 0.0673483 | 0.13933 | 0.212408 | 0.142196 |
| M589Q | F585I | 0.502054 | 0.585854 | 0.669046 | 0.596272 |
| M589Q | S586T | 0.29428 | 0.366017 | 0.426458 | 0.368788 |
| M589Q | A587P | 0.355792 | 0.404533 | 0.446143 | 0.410554 |
| M589Q | H588N | 0.539006 | 0.582224 | 0.623371 | 0.585245 |
| M589Q | N588P | -0.214526 | -0.168297 | -0.121626 | -0.169783 |
| M589Q | L588N | 0.757063 | 0.787239 | 0.817217 | 0.791389 |
| M589Q | A588N | 0.423492 | 0.479945 | 0.525762 | 0.481975 |
| M589Q | H588P | -0.140508 | -0.0813242 | -0.0190897 | -0.0811569 |
| M589Q | H588L | 0.62041 | 0.657049 | 0.686359 | 0.659036 |
| M589Q | A588H | 0.888364 | 0.910151 | 0.926381 | 0.91563 |
| M589Q | L588P | -0.159805 | -0.107828 | -0.0562832 | -0.108894 |
| M589Q | A588P | -0.161762 | -0.0932533 | -0.0222508 | -0.0925286 |
| M589Q | A588L | 0.610999 | 0.650268 | 0.68451 | 0.65374 |
| M589Q | M589T | 1 | 1 | 1 | 1 |
| M589Q | M589Q | 1 | 1 | 1 | 1 |
| M589Q | A589M | 1 | 1 | 1 | 1 |
| M589Q | Q589T | 1 | 1 | 1 | 1 |
| M589Q | A589T | 1 | 1 | 1 | 1 |
| M589Q | A589Q | 1 | 1 | 1 | 1 |
| A589M | F583H | 0.620828 | 0.639578 | 0.657849 | 0.643421 |
| A589M | L585W | 0.777034 | 0.806384 | 0.830731 | 0.809089 |
| A589M | F585W | 0.677662 | 0.716196 | 0.744622 | 0.719383 |
| A589M | I585W | 0.51177 | 0.544357 | 0.573395 | 0.546935 |
| A589M | F585L | 0.682592 | 0.702334 | 0.72173 | 0.707208 |
| A589M | I585L | 0.483193 | 0.50437 | 0.524455 | 0.507441 |
| A589M | F585I | 0.87299 | 0.890156 | 0.905071 | 0.895888 |
| A589M | S586T | 0.694659 | 0.712919 | 0.729043 | 0.717051 |
| A589M | A587P | 0.774798 | 0.789417 | 0.802622 | 0.79411 |
| A589M | H588N | 0.893239 | 0.910023 | 0.923302 | 0.911445 |
| A589M | N588P | 0.197911 | 0.264884 | 0.323634 | 0.26732 |
| A589M | L588N | 0.910349 | 0.918635 | 0.92615 | 0.920393 |
| A589M | A588N | 0.814431 | 0.834317 | 0.852222 | 0.835999 |
| A589M | H588P | 0.196481 | 0.256983 | 0.310352 | 0.259128 |
| A589M | H588L | 0.920937 | 0.935692 | 0.947719 | 0.93737 |
| A589M | A588H | 0.930429 | 0.942942 | 0.952734 | 0.945349 |
| A589M | L588P | 0.242241 | 0.306007 | 0.363388 | 0.309128 |
| A589M | A588P | 0.20626 | 0.264019 | 0.315201 | 0.267348 |
| A589M | A588L | 0.889554 | 0.904615 | 0.916889 | 0.906003 |
| A589M | M589T | 1 | 1 | 1 | 1 |
| A589M | M589Q | 1 | 1 | 1 | 1 |
| A589M | A589M | 1 | 1 | 1 | 1 |
| A589M | Q589T | 1 | 1 | 1 | 1 |
| A589M | A589T | 1 | 1 | 1 | 1 |
| A589M | A589Q | 1 | 1 | 1 | 1 |
| Q589T | F583H | 0.597285 | 0.615061 | 0.63193 | 0.617503 |
| Q589T | L585W | 0.438987 | 0.453116 | 0.467275 | 0.454343 |
| Q589T | F585W | 0.529435 | 0.553269 | 0.574216 | 0.556833 |
| Q589T | I585W | 0.272024 | 0.295169 | 0.31707 | 0.296767 |
| Q589T | F585L | 0.790603 | 0.810881 | 0.830013 | 0.813257 |
| Q589T | I585L | 0.719291 | 0.739849 | 0.760291 | 0.742441 |
| Q589T | F585I | 0.777364 | 0.800068 | 0.821324 | 0.804513 |
| Q589T | S586T | 0.624019 | 0.636605 | 0.649058 | 0.639424 |
| Q589T | A587P | 0.802956 | 0.814153 | 0.82618 | 0.817933 |
| Q589T | H588N | 0.855997 | 0.867725 | 0.877767 | 0.868739 |
| Q589T | N588P | 0.0525764 | 0.106028 | 0.167998 | 0.109564 |
| Q589T | L588N | 0.942627 | 0.949059 | 0.953995 | 0.950424 |
| Q589T | A588N | 0.77013 | 0.789992 | 0.807899 | 0.790866 |
| Q589T | H588P | 0.196399 | 0.24484 | 0.299509 | 0.247488 |
| Q589T | H588L | 0.913361 | 0.922015 | 0.928741 | 0.923086 |
| Q589T | A588H | 0.938762 | 0.950138 | 0.960523 | 0.95204 |
| Q589T | L588P | 0.0311872 | 0.0855178 | 0.143456 | 0.0868968 |
| Q589T | A588P | 0.29697 | 0.345975 | 0.396535 | 0.349837 |
| Q589T | A588L | 0.828309 | 0.846433 | 0.861905 | 0.847347 |
| Q589T | M589T | 1 | 1 | 1 | 1 |
| Q589T | M589Q | 1 | 1 | 1 | 1 |
| Q589T | A589M | 1 | 1 | 1 | 1 |
| Q589T | Q589T | 1 | 1 | 1 | 1 |
| Q589T | A589T | 1 | 1 | 1 | 1 |
| Q589T | A589Q | 1 | 1 | 1 | 1 |
| A589T | F583H | 0.218973 | 0.267318 | 0.320835 | 0.273015 |
| A589T | L585W | -0.466912 | -0.400761 | -0.334711 | -0.404836 |
| A589T | F585W | 0.173136 | 0.271737 | 0.368094 | 0.282318 |
| A589T | I585W | -0.0826207 | 0.0444774 | 0.157416 | 0.0433943 |
| A589T | F585L | 0.161457 | 0.221944 | 0.276948 | 0.225876 |
| A589T | I585L | 0.365603 | 0.422121 | 0.476749 | 0.430058 |
| A589T | F585I | 0.16008 | 0.287982 | 0.406716 | 0.300457 |
| A589T | S586T | 0.161759 | 0.219948 | 0.278359 | 0.224516 |
| A589T | A587P | 0.573639 | 0.629198 | 0.6743 | 0.640881 |
| A589T | H588N | 0.561835 | 0.6262 | 0.675585 | 0.632318 |
| A589T | N588P | 0.013092 | 0.0796778 | 0.150175 | 0.0818501 |
| A589T | L588N | 0.708985 | 0.744835 | 0.770778 | 0.753502 |
| A589T | A588N | 0.549862 | 0.599588 | 0.642558 | 0.60824 |
| A589T | H588P | -0.0347897 | 0.0301399 | 0.102947 | 0.0301491 |
| A589T | H588L | 0.627012 | 0.671714 | 0.709238 | 0.679601 |
| A589T | A588H | 0.714481 | 0.768505 | 0.819705 | 0.779044 |
| A589T | L588P | 0.0145005 | 0.0767062 | 0.146671 | 0.0791503 |
| A589T | A588P | 0.0839389 | 0.151211 | 0.217324 | 0.154786 |
| A589T | A588L | 0.596823 | 0.651984 | 0.695319 | 0.658814 |
| A589T | M589T | 1 | 1 | 1 | 1 |
| A589T | M589Q | 1 | 1 | 1 | 1 |
| A589T | A589M | 1 | 1 | 1 | 1 |
| A589T | Q589T | 1 | 1 | 1 | 1 |
| A589T | A589T | 1 | 1 | 1 | 1 |
| A589T | A589Q | 1 | 1 | 1 | 1 |
| A589Q | F583H | 0.566362 | 0.594292 | 0.622859 | 0.599141 |
| A589Q | L585W | 0.842518 | 0.87691 | 0.900334 | 0.880724 |
| A589Q | F585W | 0.757333 | 0.784199 | 0.80782 | 0.789586 |
| A589Q | I585W | 0.511474 | 0.550548 | 0.584331 | 0.553989 |
| A589Q | F585L | 0.647221 | 0.678017 | 0.704434 | 0.682002 |
| A589Q | I585L | 0.34506 | 0.382892 | 0.417525 | 0.385933 |
| A589Q | F585I | 0.778609 | 0.817476 | 0.850239 | 0.824274 |
| A589Q | S586T | 0.527521 | 0.555718 | 0.578013 | 0.559076 |
| A589Q | A587P | 0.779712 | 0.799686 | 0.819184 | 0.805849 |
| A589Q | H588N | 0.850867 | 0.875346 | 0.896006 | 0.876859 |
| A589Q | N588P | 0.368269 | 0.427761 | 0.482045 | 0.433317 |
| A589Q | L588N | 0.88703 | 0.89749 | 0.906085 | 0.8993 |
| A589Q | A588N | 0.759905 | 0.788345 | 0.812009 | 0.790563 |
| A589Q | H588P | 0.334314 | 0.394742 | 0.452035 | 0.400742 |
| A589Q | H588L | 0.926399 | 0.942833 | 0.956197 | 0.945271 |
| A589Q | A588H | 0.916844 | 0.932468 | 0.946369 | 0.937113 |
| A589Q | L588P | 0.391347 | 0.447345 | 0.495941 | 0.452695 |
| A589Q | A588P | 0.295553 | 0.356449 | 0.414503 | 0.361675 |
| A589Q | A588L | 0.873165 | 0.89395 | 0.911214 | 0.896538 |
| A589Q | M589T | 1 | 1 | 1 | 1 |
| A589Q | M589Q | 1 | 1 | 1 | 1 |
| A589Q | A589M | 1 | 1 | 1 | 1 |
| A589Q | Q589T | 1 | 1 | 1 | 1 |
| A589Q | A589T | 1 | 1 | 1 | 1 |
| A589Q | A589Q | 1 | 1 | 1 | 1 |

**Table S4_2.** The average epistatic effect *γA*_*i*_→ (eq. S1_9) of a mutation occurring on any background calculated over 1,000 posterior samples. Shown are the 2.5%, 50%, 97.5% confidence intervals and the corresponding value calculated from the median growth rates.

| $A_i$ | 2.5%C.I. | 50%C.I. | 97.5%C.I. | Main text |
| --- | --- | --- | --- | --- |
| F583 | 0.645879 | 0.655648 | 0.665972 | 0.659633 |
| H583 | 0.645879 | 0.655648 | 0.665972 | 0.659633 |
| W585 | 0.509098 | 0.519484 | 0.529889 | 0.523148 |
| L585 | 0.494295 | 0.504828 | 0.515661 | 0.509236 |
| F585 | 0.480383 | 0.493608 | 0.506724 | 0.500503 |
| T585 | 0.524727 | 0.537596 | 0.549722 | 0.542512 |
| S586 | 0.574784 | 0.58707 | 0.598905 | 0.591946 |
| T586 | 0.574784 | 0.58707 | 0.598905 | 0.591946 |
| A587 | 0.470531 | 0.484783 | 0.498591 | 0.491348 |
| P587 | 0.470531 | 0.484783 | 0.498591 | 0.491348 |
| N588 | 0.753967 | 0.760283 | 0.766723 | 0.763474 |
| H588 | 0.714414 | 0.722585 | 0.72996 | 0.726474 |
| P588 | 0.790154 | 0.79631 | 0.802035 | 0.799044 |
| L588 | 0.78632 | 0.792616 | 0.798058 | 0.795508 |
| A588 | 0.69515 | 0.702644 | 0.709753 | 0.706728 |
| M589 | 0.625289 | 0.637087 | 0.648515 | 0.640628 |
| T589 | 0.601966 | 0.610839 | 0.620123 | 0.614661 |
| Q589 | 0.614382 | 0.625208 | 0.635169 | 0.629363 |
| A589 | 0.59604 | 0.608972 | 0.621588 | 0.614412 |

**Table S4_3.** The average epistatic effect *γ*→*A*_*j*_ (eq. S1_13) of a background on any new mutation calculated over 1,000 posterior samples. Shown are the 2.5%, 50%, 97.5% confidence intervals and the corresponding value calculated from the median growth rates. In accordance with our results from the main text the average epistatic effect of a background on any new mutation is usually small, except for background P588*, which shows a strong magnitude effect.

| $A_j$ | 2.5%C.I. | 50%C.I. | 97.5%C.I. | Main text |
| --- | --- | --- | --- | --- |
| 583F | 0.643547 | 0.654621 | 0.665255 | 0.658462 |
| 583H | 0.643547 | 0.654621 | 0.665255 | 0.658462 |
| 585W | 0.623883 | 0.633104 | 0.641805 | 0.636432 |
| 585L | 0.733728 | 0.740188 | 0.746721 | 0.743645 |
| 585F | 0.777099 | 0.78529 | 0.792513 | 0.7899 |
| 585T | 0.659718 | 0.67033 | 0.680289 | 0.674306 |
| 586S | 0.709667 | 0.720978 | 0.730423 | 0.724845 |
| 586T | 0.709667 | 0.720978 | 0.730423 | 0.724845 |
| 587A | 0.774192 | 0.782478 | 0.789768 | 0.78691 |
| 587P | 0.774192 | 0.782478 | 0.789768 | 0.78691 |
| 588N | 0.60486 | 0.614512 | 0.62454 | 0.618562 |
| 588H | 0.673788 | 0.682904 | 0.692253 | 0.687369 |
| 588P | 0.14836 | 0.172296 | 0.194468 | 0.17391 |
| 588L | 0.650123 | 0.659805 | 0.669129 | 0.663803 |
| 588A | 0.658737 | 0.668566 | 0.677671 | 0.672761 |
| 589M | 0.601843 | 0.612576 | 0.622367 | 0.615955 |
| 589T | 0.587559 | 0.599117 | 0.61039 | 0.602947 |
| 589Q | 0.706356 | 0.714376 | 0.722292 | 0.717983 |
| 589A | 0.667143 | 0.678345 | 0.689501 | 0.683378 |

**Table S4_4.** Observed decay of landscape-wide epistasis measure *γ*_*d*_ (eq. S1_15) with mutational distance *d* calculated over 1,00 posterior samples. Shown are the 2.5%, 50%, 97.5% confidence intervals and the corresponding value calculated from the median growth rates. Result based on the posterior samples support our finding from the main text, i.e., that the general topography of the fitness landscape resembles that of a RMF landscape with intermediate ruggedness

| Mutational distance $d$ | 2.5%C.I. | 50%C.I. | 97.5%C.I. | Main text |
| --- | --- | --- | --- | --- |
| 0 | 1 | 1 | 1 | 1 |
| 1 | 0.656709 | 0.663385 | 0.668302 | 0.667961943 |
| 2 | 0.565145 | 0.571478 | 0.576907 | 0.575378659 |
| 3 | 0.529712 | 0.535176 | 0.540732 | 0.539030991 |
| 4 | 0.512834 | 0.517746 | 0.522954 | 0.521536863 |
| 5 | 0.509929 | 0.51507 | 0.519867 | 0.518957681 |

**Table S4_5.** Roughness-to-slope ratio calculated over 1,000 posterior samples. Shown are the 2.5%, 50%, 97.5% confidence intervals and the corresponding value calculated from the median growth rates. Variation between measures is very low, all supporting a fitness landscape consisting of a mixture of a random HoC component and an additive component (i.e., a RMF landscape).

| 2.5%C.I. | 50%C.I. | 97.5%C.I. | Main text |
| --- | --- | --- | --- |
| 0.910872 | 0.92082 | 0.930658 | 0.915688 |

**Table S4_6.** The type and frequency of the global optimum sequence for the 1,000 posterior fitness landscapes. Colored cells represent the mutant that was found to be the global optimum in the median growth rate fitness landscape (i.e., in the main text).

| Sequence | Frequency |
| --- | --- |
| QFGWSPLAE | 3 |
| QFGFSALTE | 1 |
| QFGLSPLAE | 996 |

**Table S4_7.**
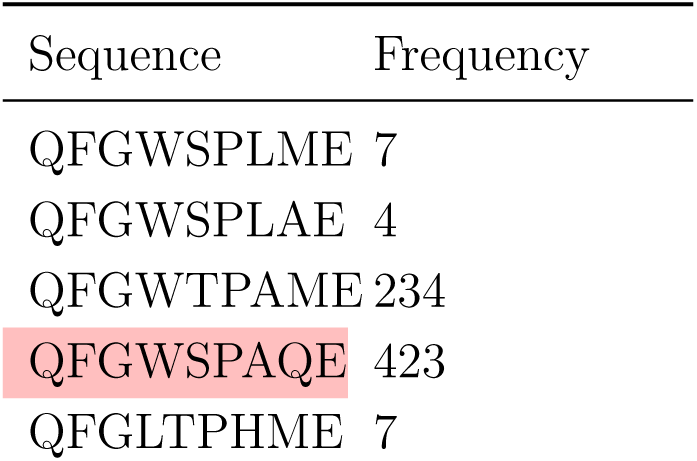
The type and frequency of the local optima sequences for the 1,000 posterior fitness landscapes. Colored cells represent those mutants that were found to be the local optima in the median growth rate fitness landscape (i.e., in the main text).

**Table S4_8.**
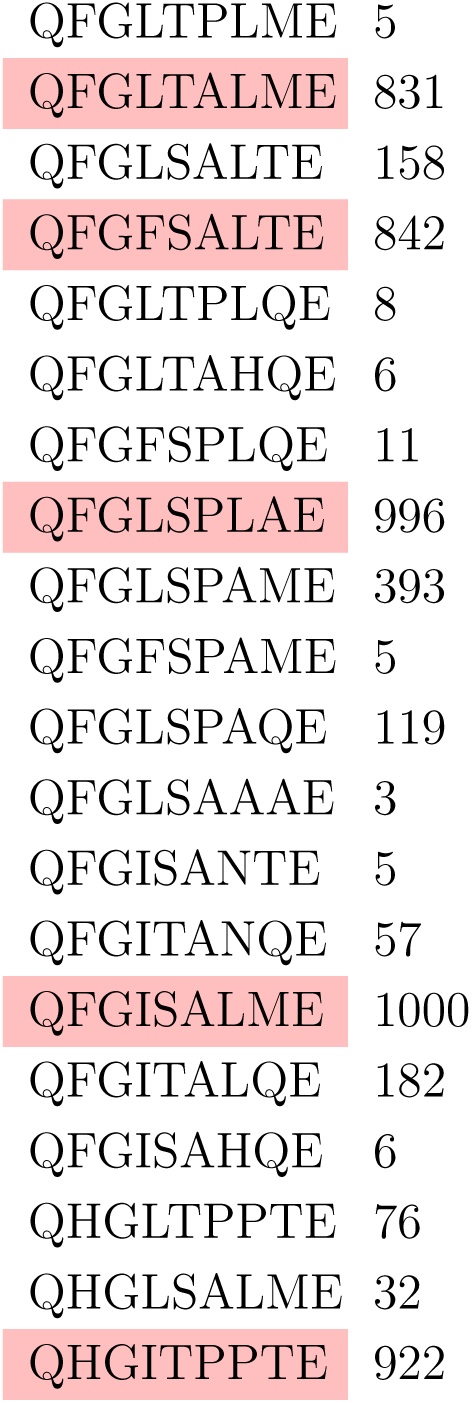
The number of local optima for the 1,000 posterior fitness landscapes. The colored cell represents the number of local optima that was found in the median growth rate fitness landscape (i.e., in the main text).

